# KIF1A-mediated trafficking is required for neuronal autophagy in human neurons

**DOI:** 10.64898/2026.07.22.740140

**Authors:** Carris Borland, Jacob Popolow, Erika L. F. Holzbaur

## Abstract

Mutations in the molecular motor protein KIF1A result in a spectrum of neurodevelopmental and neurodegenerative disorders termed KIF1A-Associated Neurological Disorder (KAND). KIF1A mutations variably disrupt synaptic vesicle trafficking, but the effects of KIF1A mutations on other trafficking pathways remain unexplored. Autophagy is a conserved pathway required for neuronal homeostasis. We investigated the role of KIF1A in autophagy using gene-edited human IPSC-derived neurons. KIF1A loss inhibited the trafficking of ATG9, a transmembrane lipid scramblase necessary for autophagosome biogenesis. This deficit significantly reduced autophagosome biogenesis and the density of axonal autophagosomes. KIF1A loss also depleted lysosomes from the axon, inhibiting autophagosome maturation. In neurons gene-edited to heterozygously express a pathogenic variant linked to a Rett-like syndrome in KAND patients, we also noted significant deficits in autophagy and lysosomal trafficking. Together, these results suggest that KIF1A-mediated transport is critical to neuronal autophagy and that deficits in autophagy may contribute to pathogenesis in KAND.

**GRAPHICAL ABSTRACT:** 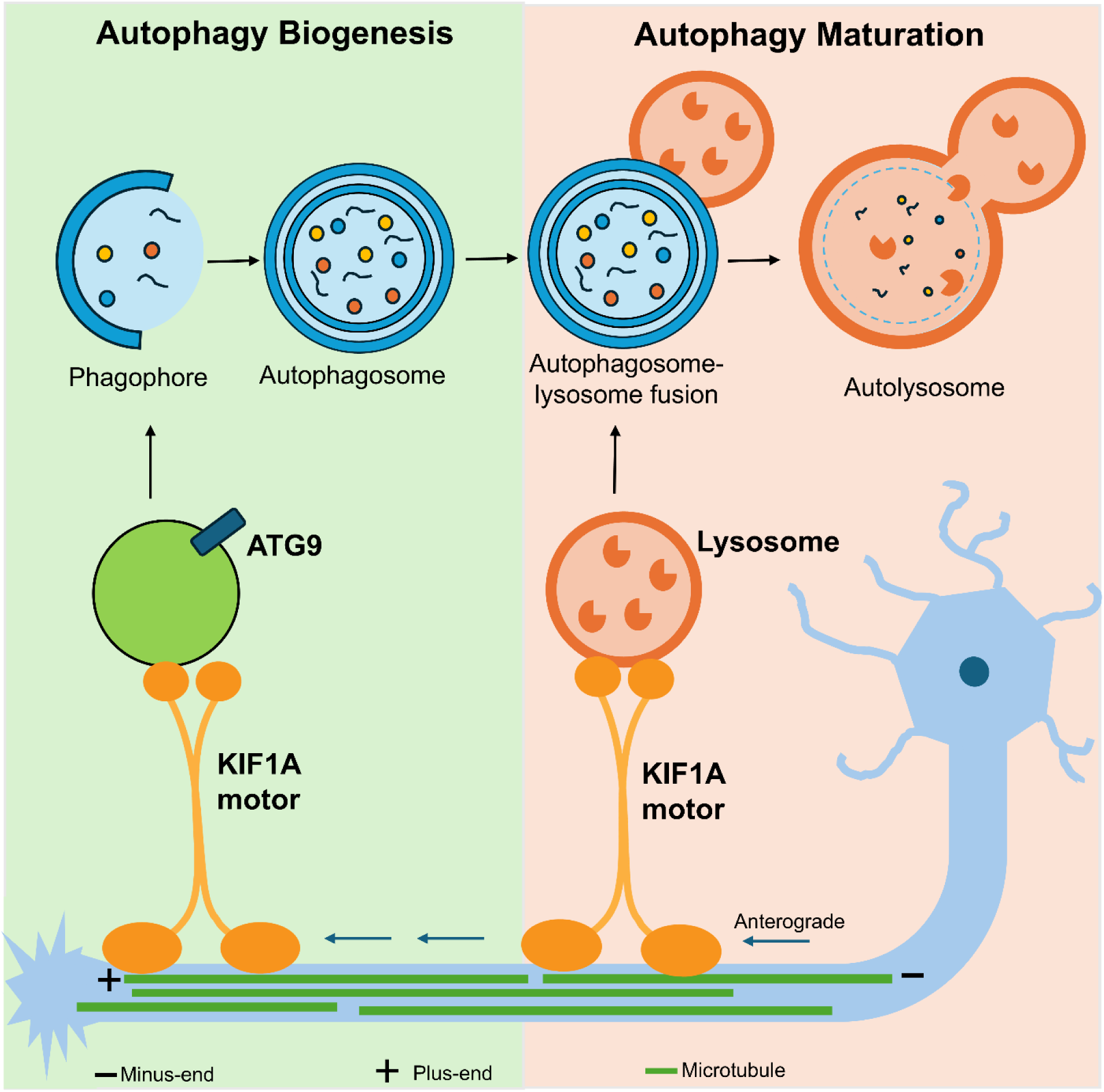

## INTRODUCTION

Neurons rely on axonal transport for the targeted and efficient delivery of intracellular components. Molecular motor proteins drive the transport of organelles, vesicles, RNA granules, and proteins along the polarized axonal microtubule array, precisely delivering essential materials to presynaptic sites and the axon terminal. Kinesin motors drive anterograde trafficking along the axon, while cytoplasmic dynein and its essential activator dynactin drive retrograde transport back to the soma.^1^ Highlighting the importance of this transport, both neurodegenerative and neurodevelopmental disorders have been linked to deficits in axonal transport.^2, 3^ KIF1A is a well-characterized motor in the kinesin-3 family, which was initially identified in *C. elegans* as a motor essential for the transport of synaptic vesicle precursors (SVPs).^4^ In mammalian neurons, KIF1A drives the transport of both SVPs and dense core vesicles (DCVs).^5,6^ KIF1A-mediated trafficking is fundamental for neuronal viability, robust synaptic transmission, synaptogenesis, and learning and memory.^7,8,9^

Structurally, KIF1A has an N-terminal motor domain followed by a neck domain and a forkhead-associated (FHA) domain.^10^ The motor is found in the cytosol in a monomeric, auto-inhibited state, but dimerizes and becomes active upon cargo binding.^11,12^ KIF1A is a highly processive microtubule motor, due to a lysine-rich K-loop in the motor domain that enhances interaction with the microtubule, and a short neck-linker that encourages tight motor-head stepping coordination.^13,14^ Furthermore, in the presence of hindering load, KIF1A can rapidly re-engage with the microtubule following detachment, enabling efficient motility and navigation of obstacles.^15,16^ These unique motile properties give KIF1A the ability to engage in fast, highly processive long-distance transport of SVPs and dense core vesicles along the axon.

Pathogenic KIF1A variants result in a spectrum of neurodevelopmental and neurodegenerative deficits collectively called KIF1A-associated neurological disorder (KAND). Symptoms can include spastic paraplegia, microcephaly, encephalopathy, intellectual disability, autism, autonomic and peripheral neuropathy, optic nerve atrophy, cerebral and cerebellar atrophy, and epilepsy.^17,18,19,20^ Additionally, mutations in KIF1A have been linked to Amyotrophic lateral sclerosis (ALS).^21,22^ Most mutations in KIF1A are heterozygous *de novo* variants and are distributed throughout the sequence, with the highest frequency occurring in the motor domain.^15,19^ Clinical severity largely depends on the type of mutation and its location within the gene.^23,24^ Both loss-of-function and gain-of-function mutations have been characterized using in vitro and cellular approaches, as well as in model organisms.^15,23,24,25,26,27^ There has been strong progress in elucidating the molecular phenotypes of specific KAND-associated mutations and how each mutation impacts KIF1A functionality. Research on the cellular consequences of KAND variants has centered primarily on synaptic trafficking and function. However, KIF1A is known to transport other cargos, such as lysosomes.^28,29^ Work from *C. elegans* has shown that KIF1A transports ATG9-containing vesicles.^30^ ATG9 vesicles are required for autophagosome biogenesis, interacting with ATG2 to provide membrane lipids to forming autophagosomes.^31,32^ These observations suggest that mutations in KIF1A may detrimentally affect the degradative capacity of neuronal axons. Autophagy is a vital homeostatic pathway indispensable for neuronal health. In neurons, autophagy is temporally and spatially regulated, occurring robustly under basal conditions to remove aging mitochondria and synaptic vesicles from presynaptic sites in the distal axon.^33,34,35^ Cargoes are sequestered within a double-membrane vesicle that is recruited to axonal microtubules and undergoes rapid transport to the cell soma, driven by cytoplasmic dynein and dynactin. Autophagosomes fuse with lysosomes en route, maturing to degradation-competent organelles.^34^ The autophagy pathway enables efficient turnover of proteins and organelles, neuronal development, regulation of learning and memory, and the maintenance of synaptic integrity.^36,37^ Furthermore, defects in autophagy are linked to many neurodegenerative and neurodevelopmental diseases,^38,39,40^ raising the possibility that autophagic deficits caused by pathogenic KIF1A mutations may contribute to the clinical spectrum seen in KAND patients.

Here, we investigate the impact of KIF1A-mediated trafficking on axonal autophagy using human iPSC-derived neurons gene-edited to express an early stop codon in KIF1A, C92*, identified as a pathogenic loss-of-function mutation causal for KAND^25,41^ (**Figure 1A**). Both PCR analysis and immunoblotting indicate this is a null mutation when homozygous.^25^ We find that loss of KIF1A in neurons differentiated from a homozygous C92* null line impacts both the formation and the maturation of autophagosomes in human neurons. Loss of KIF1A results in ATG9 mislocalization, reduced autophagosome biogenesis, reduced density of both autophagosomes and lysosomes along the axon, and inhibition of autophagosome maturation. We also examined iPSC-derived neurons gene-edited to be heterozygous for the C92* early truncation mutation (C92*Het). KIF1A expression is significantly reduced in heterozygous neurons, and strikingly, we find significant deficits in autophagosome biogenesis and maturation, indicating that KIF1A is not haplo-sufficient for autophagy in human neurons. Together, these findings demonstrate an essential role for KIF1A in neuronal autophagy and suggest that autophagic deficits may contribute to the KAND phenotype in affected individuals.

**Figure 1:**
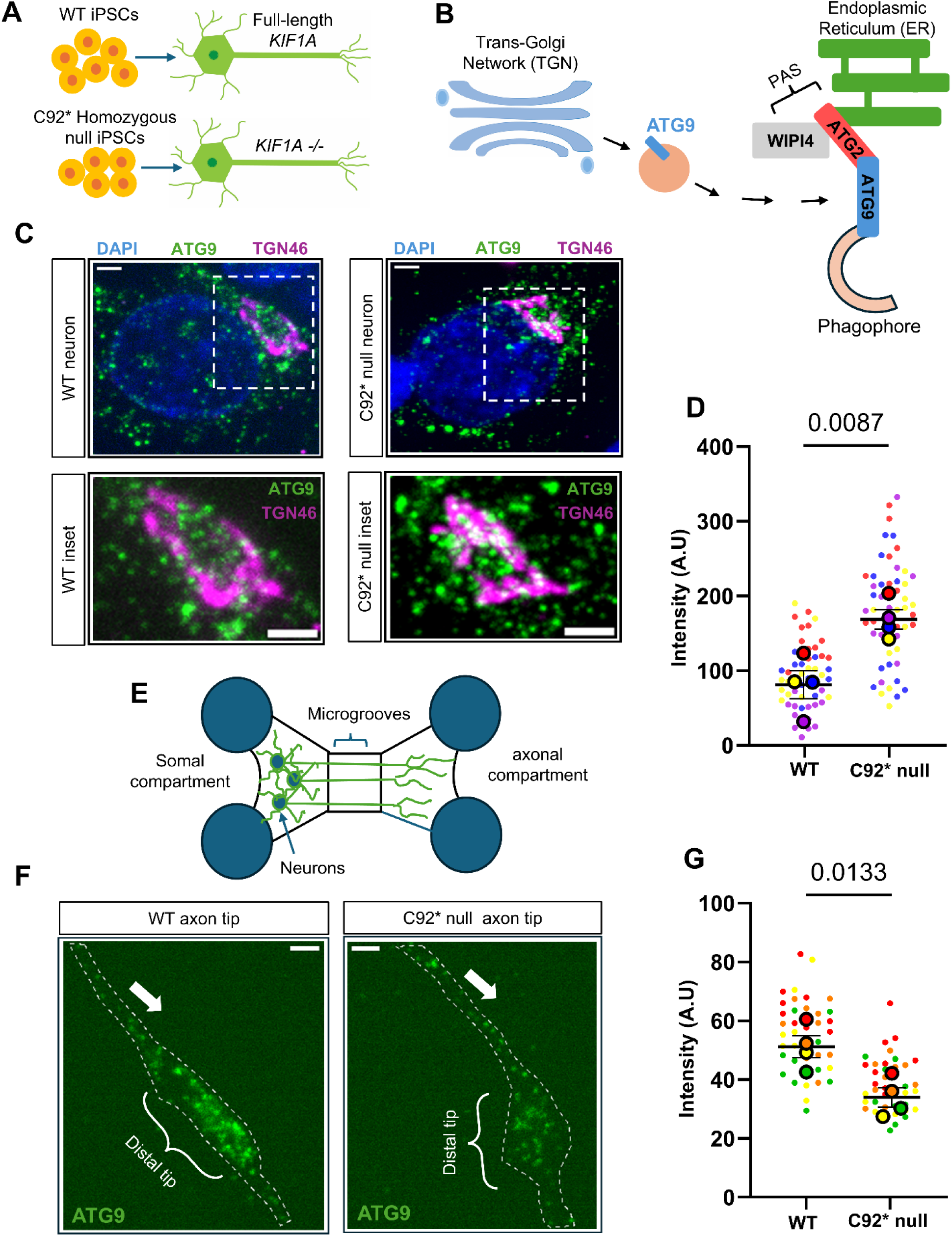
ATG9 localization is disrupted in the absence of KIF1A. **A)** WT iPSCs and iPSCs gene-edited to endogenously express the C92* homozygous truncating variant were differentiated to cortical-like glutamatergic neurons. **B)** Schematic showing the role of ATG9 in phagophore expansion. ATG9-containing vesicles originate from the trans-Golgi network (TGN) and are trafficked to the phagophore assembly site (PAS). The ATG9 vesicle is the “seed” membrane for the growing phagophore, docking at the endoplasmic reticulum (ER) via the ATG2-WIPI4 complex. ATG2 channels lipids from the ER, then ATG9 further facilitates lipid transfer by acting as a lipid scramblase, thus leading to phagophore membrane expansion. **C)** Representative images of ATG9 localization in the soma of WT and C92* null neuron. White dotted box signifies the areas of the TGN shown enlarged in the insets below. DAPI (blue), ATG9 (green), TGN46 (magenta). Scale bars, 2µm. **D)** Mean grey value of ATG9 intensity confined to the TGN. Plot shows mean ± standard deviation of experimental replicates; n = 52 neurons from 4 independent experiments. Small dots represent individual cells; large dots represent the average for each replicate. Each color represents a biological replicate. WT average = 93, C92* null average= 179. P-value determined by unpaired t-test. **E)** Schematic showing neuronal culture within a microfluidic chamber enabling physical separation of soma and axons. Microscopic microgrooves are designed to restrict cell bodies to one compartment (somal compartment) while promoting neuronal axons to grow into the adjacent compartment (axonal compartment). **F)** Representative images of ATG9 localization at the distal tip of WT and C92* null axons. Neurons were cultured in microfluidic chambers to physically isolate axons and stained with ATG9 (green) and neurofilament-H (NFH; not shown) antibodies. White dotted lines signify the outline of the distal tips. Arrows point toward the distal end of the axon. Scale bars 5µm. **G)** Mean grey value of ATG9 intensity confined to the distal tip. Plot shows mean ± standard deviation of experimental replicates; n= 40 neurons from 4 independent experiments. Small dots represent individual axons; large dots represent the average for each replicate. Each color represents 1 replicate. WT average= 55, C92* null average= 38. P-value determined by unpaired t-test.

## RESULTS

### Loss of KIF1A disrupts ATG9 localization in human neurons

Autophagy biogenesis in mammalian neurons begins with the de novo formation of a cup-shaped double-membrane precursor, called the phagophore, at endoplasmic reticulum (ER) contact sites called phagophore assembly sites (PAS) in the distal axon^42,43,44^ The phagophore matures into an autophagosome by acquiring lipids from the ER for elongation and expansion. ATG9 is a transmembrane protein that is one of the first to be recruited to the PAS, and thought to act as a “seed” for the growing phagophore^45^, interacting with other components of the autophagy machinery such as the ATG2-WIPI4 complex to facilitate incorporation of lipids into the growing phagophore^45,46^ (**Figure 1B**). ATG9, a homotrimer with a central hydrophilic pore, is a scramblase that redistributes lipids transferred from the ER by the bridge-like lipid transfer protein ATG2 by “flipping” them from one leaflet to another. ATG9’s scramblase activity is essential for autophagosome formation, as ATG9 knockout cells or cells expressing mutations in the ATG9 hydrophilic core exhibit defective phagophore expansion and autophagy formation.^45,47,48,31^ ATG9-vesicles are Golgi-derived and are trafficked to the distal axon for autophagy biogenesis.^49,50^ In C. elegans, ATG9 vesicles are trafficked to autophagy initiation sites in the axon by the KIF1A motor.^30^ To determine whether KIF1A is required for ATG9 distribution to sites of autophagy biogenesis in human neurons, we employed immunocytochemistry to detect steady state levels of endogenous ATG9 in the soma and distal axons of WT and p.C92* null neurons (**Figure 1C**). ATG9 distribution was significantly altered in neurons lacking KIF1A. Intensity measurements reveal that KIF1A null neurons exhibited increased confinement of ATG9 to the trans-Golgi network (TGN) within the soma as compared to WT neurons (**Figure 1C-1D**). To investigate ATG9 enrichment in the distal axon, we cultured WT and p.C92* null neurons in microfluidic chambers to physically separate axons and then stained for ATG9 (**Figure 1E**). Consistent with aberrant somal retention, we observed significantly decreased ATG9 levels in the distal axon in KIF1A-null neurons compared to WT neurons (**Figure 1F-1G**). These observations indicate that KIF1A is required for normal ATG9 trafficking to the distal axon of human neurons, consistent with previous observations in *C. elegans*, and that loss of KIF1A results in the distal depletion of ATG9-containing vesicles.

### Loss of KIF1A impairs distal autophagosome density and autophagosome flux along the axon

Next, we assessed the impact of KIF1A-mediated ATG9 mislocalization on autophagosome density in the distal axon by culturing WT and C92* null neurons in microfluidic chambers, using live-cell imaging to visualize endogenous autophagosomes with DAPRED. DAPRED is a small fluorescent probe that is incorporated into the membrane of a forming autophagosome^51^ **(Figure 2A)**. Mammalian autophagosomes typically range between 0.5-1.5µm in diameter,^52,53,54,55^ therefore, we counted the density of DAPRED-positive autophagosomes 0.5-1.5µm in diameter in the distal axon, defined as the region within 100 µm of the axon tip. In p.C92* null axons, there were only half as many distal DAPRED- labeled autophagosomes as compared to WT axons **(Figure 2B-E)**. Next, we asked if this decrease in the density of distal autophagosomes impacted the number of organelles trafficking along the axon. We transfected WT and p.C92* null neurons with LC3B tagged with the red fluorescent protein mScarlet and used live-cell imaging to capture time-lapse videos. The videos were used to generate kymographs to track autophagosome vesicle movement over time **(Figure 2F).** We saw significantly fewer autophagosomes traveling along KIF1A-null axons compared to WT **(Figure 2G-H)**. Both the reduced density of distal autophagosomes and the corresponding reduction in the trafficking of autophagosome suggest there is an inhibition of autophagy biogenesis in KIF1A- null neurons.

**Figure 2:**
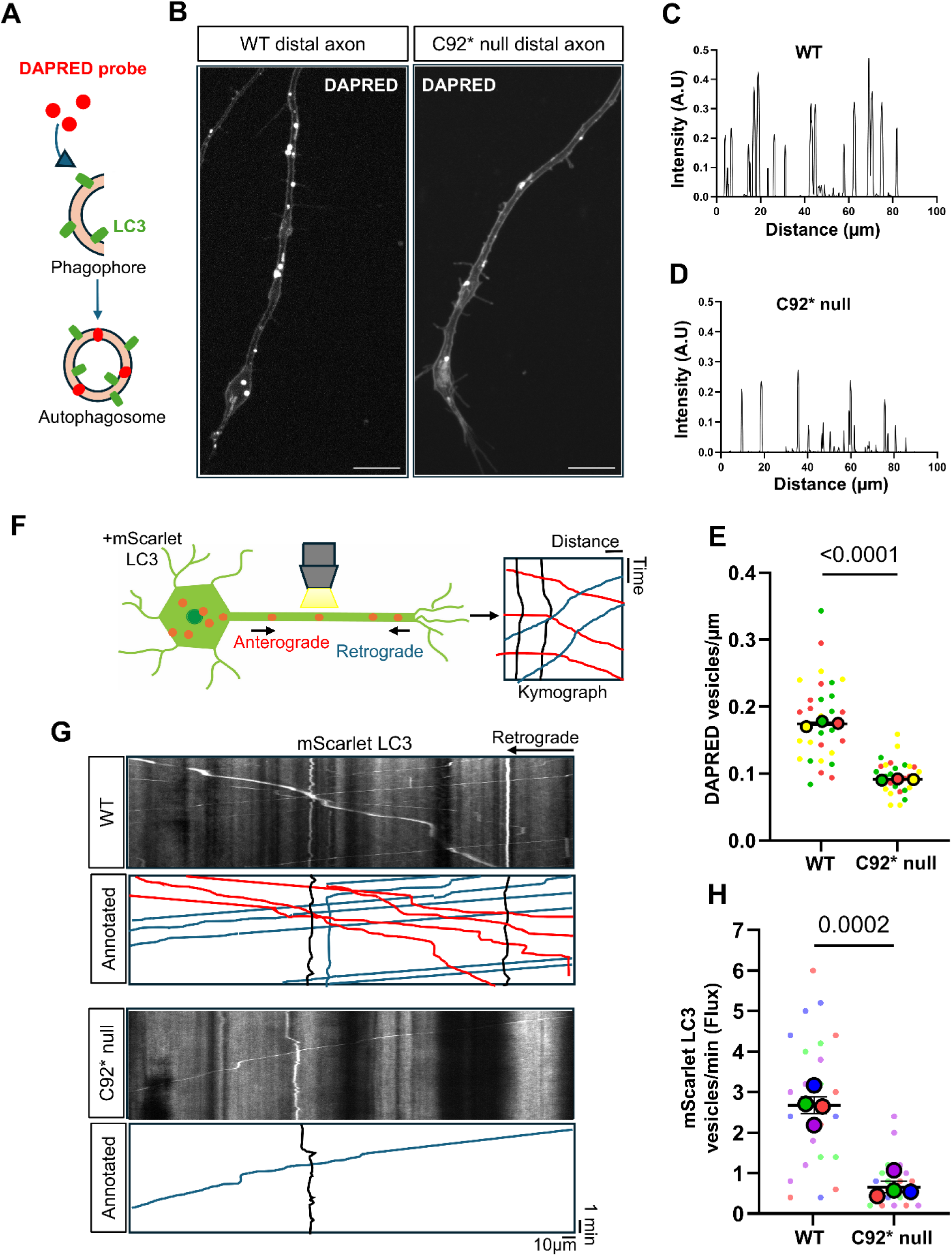
KIF1A loss impacts autophagosome density and flux. **A)** DAPRED is a cell-permeable fluorescent molecule that is incorporated into the membrane of a forming autophagosome (phagophore) and allows for real-time monitoring of endogenous autophagosomes in live cells. **B)** DAPRED puncta in WT and C92* null distal axons. Neurons were cultured in microfluidic chambers, and the distal axons were imaged at DIV7. The distal axon region was defined as 100µm from the very tip of the axon. Puncta diameter size between 0.5-1.5µm were segmented and counted. Scale bars 10µm. **C)** Intensity linescan for the WT distal axon shown in 2B stained with DAPRED. Y-axis is mean grey value; x-axis is distance in microns. **D)** Intensity linescan for a C92* null distal axon shown in 2B stained with DAPRED. Y-axis is mean grey value; x-axis is distance in microns. **E)** Superplot showing the number of DAPRED puncta in the distal axons of WT and C92* null axons (vesicle/µm). Plot shows mean ± standard deviation of experimental replicates; n =30 neurons for each condition from 3 independent experiments. Small dots represent individual cells; large dots represent the average for each replicate. Each color represents an individual biological replicate. WT average = 0.2 vesicles/ µm, C92* null average= 0.1 vesicles/ µm. P-value determined by unpaired t-test. **F)** Timelapse live cell imaging of autophagic vesicle flux in DIV21 WT and C92* null neurons transfected with mScarlet-LC3 plasmid. Time lapse videos were collected, and subsequent kymographs were generated to track vesicle movement over time. **G)** Kymographs with annotations showing the flux of mScarlet-LC3-positive vesicles in WT and C92* null axons. X axis is distance, Y axis is time. Time-lapse videos were recorded for 5 minutes at 5fps. Red tracks show anterograde moving LC3 vesicles; blue tracks show retrograde moving LC3 vesicles; black tracks show stalled LC3 vesicles. Arrow pointing in retrograde direction. Scale bars 10µm. **H)** Superplot showing the total LC3 vesicle flux (vesicles/min) in WT and C92* null axons. Plot shows mean ± standard deviation of experimental replicates; n =24 neurons for each condition from 4 independent experiments. Small dots represent individual cells; large dots represent the average for each replicate. Each color represents 1 replicate. WT average = 2.8 vesicles/min, C92* null average= 0.8 vesicles/min. P-value determined by unpaired t-test.

### KIF1A depletion inhibits autophagosome biogenesis

To determine whether the observed reduction in the density of distal autophagosomes is caused by a deficit in autophagosome formation, we employed live-cell imaging to measure rates of autophagosome biogenesis. Autophagosome formation follows an orderly recruitment of autophagy machinery proteins.^33,44,56^ DFCP1 (Double FYVE Domain-Containing Protein 1) is an ATPase that is recruited to PI3P-rich sites on the ER called omegosomes,^57^ making it a useful marker for the initiation of autophagosome biogenesis. The ATPase activity of DFCP1 drives the constriction and eventual release of nascent autophagosomes from the omegosome^57,58^ **(Figure 3A).** DIV21 WT and p.C92* null neurons were co-transfected with Halo-DFCP1 and mScarlet-LC3, and 15-minute videos of the distal axon tips were recorded to capture the colocalization of DFCP1 and LC3 puncta (**Figure 3B-C**). LC3 puncta that did not colocalize with DFCP1 during the video recordings were not counted as biogenesis events. Indeed, we saw significantly fewer biogenesis events occurring in C92* null axons compared to WT **(figure 3D-F**). These findings indicate that the reduced steady state distribution of autophagosomes observed in C92* null neurons is a result of reduced biogenesis in the distal axon.

**Figure 3:**
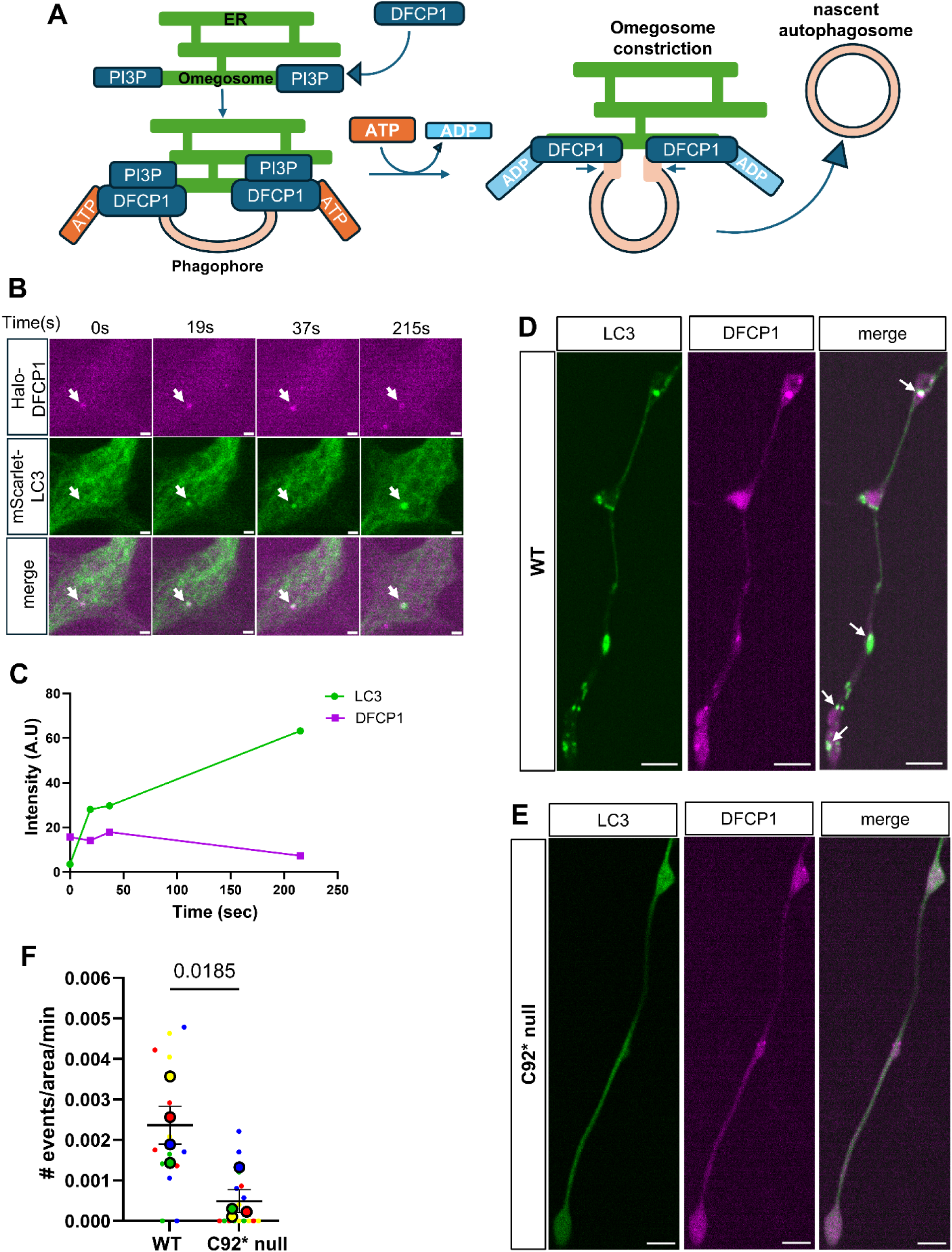
KIF1A absence result in slower autophagy biogenesis rates. **A)** Schematic showing DFCP1’s role in autophagosome biogenesis. The de novo formation and expansion of an autophagosome begins with the recruitment of DFCP1 to PI3P-enriched subdomains of the endoplasmic reticulum (ER) called omegosomes. DFCP1’s ATPase activity drives the constriction of the omegosome and eventually releases a nascent autophagosome from the ER. **B)** Time series showing the assembly and disassembly of DFCP1 as an autophagosome forms. DFCP1 is recruited to the omegosome at the start of autophagosome formation. As the autophagosome grows, DFCP1 intensity decreases, which coincides with the release of the autophagosome. 15-minute videos of the distal axons were recorded at 1fps. Only autophagosomes co-localizing with DFCP1 was considered as a biogenesis event. Scale bars 2µm. **C)** Intensity plot showing the dynamics of DFCP1 assembly and disassembly as an autophagosome grows. Y-axis is intensity (mean grey value); x-axis is time in seconds. **D)** Representative images of WT distal axon co-transfected with mScarlet LC3 (green) and Halo-DFCP1 (magenta). Arrows point to LC3 puncta that were considered biogenesis events. Scale bar 5µm. **E)** Representative images of C92*null distal axon co-transfected with mScarlet LC3 (green) and Halo-DFCP1 (magenta). Arrows point to LC3 puncta that were considered biogenesis events. Scale bar 5µm. **F)** Superplot showing the rate of biogenesis events in the distal axons of WT and C92* null axons (#events/area/min). Plot shows mean ± standard deviation of experimental replicates; n =16 neurons for each condition from 4 independent experiments. Small dots represent individual cells; large dots represent the average for each replicate. Each color represents 1 replicate. WT average = 0.002 events/area/min, C92* null average= 0.001 events/area/min. P-value determined by unpaired t-test with Welch’s correction.

### Lysosomes are depleted from KIF1A-null axons

Lysosomes are dynamic organelles involved in degradation and recycling, cell signaling and nutrient sensing.^59^ Autophagosomes must fuse with lysosomes to become functional degradative organelles; this fusion occurs in the axon during trafficking of autophagosomes toward the soma.^34^ Lysosome distribution in the axon is also motor-dependent.^60^ Both kinesin-1 and kinesin-3 motors have been implicated in this trafficking;^29,61,28,62^ the activities of these kinesin motors are regulated by the BORC complex.^63^ First, we established that KIF1A co-localizes with LAMP1-positive lysosomes **(Figure 4A-4B)** and that KIF1A co-migrates with LAMP1-positive lysosomes in axons **(Figure 4C).** We then asked whether lysosome distribution is impacted by the absence of KIF1A. Neurons at DIV21 were transfected with mNeon-LAMP1. Axons of C92* null neurons exhibited significantly lower lysosomal density compared to WT neurons **(Figure 4D-G)**. To determine whether the observed depletion of lysosomes seen in C92* null axons was accompanied by a corresponding increase in lysosomal density within the soma, WT and C92* null neurons were immunostained for endogenous levels of LAMP1. Intensity measurements revealed increased levels of LAMP1 in the perinuclear region of the soma for C92* neurons compared to WT **(Figure 4H-4J).** As neurons can increase lysosome production in response to stress, we investigated whether these changes in lysosome positioning were accompanied by increased lysosome expression. Comparisons of cell lysates from KIF1A-null and WT neurons did not indicate significant changes in levels of either LAMP1 (**Supplemental figure 1A and 1B)** or cathepsin D (**Supplemental figure 1A and 1C)**. Nor did we see significant changes in levels of phospho-TFEB, a regulator of lysosome biogenesis **(Supplemental figure 1A and 1D).**

**Figure 4:**
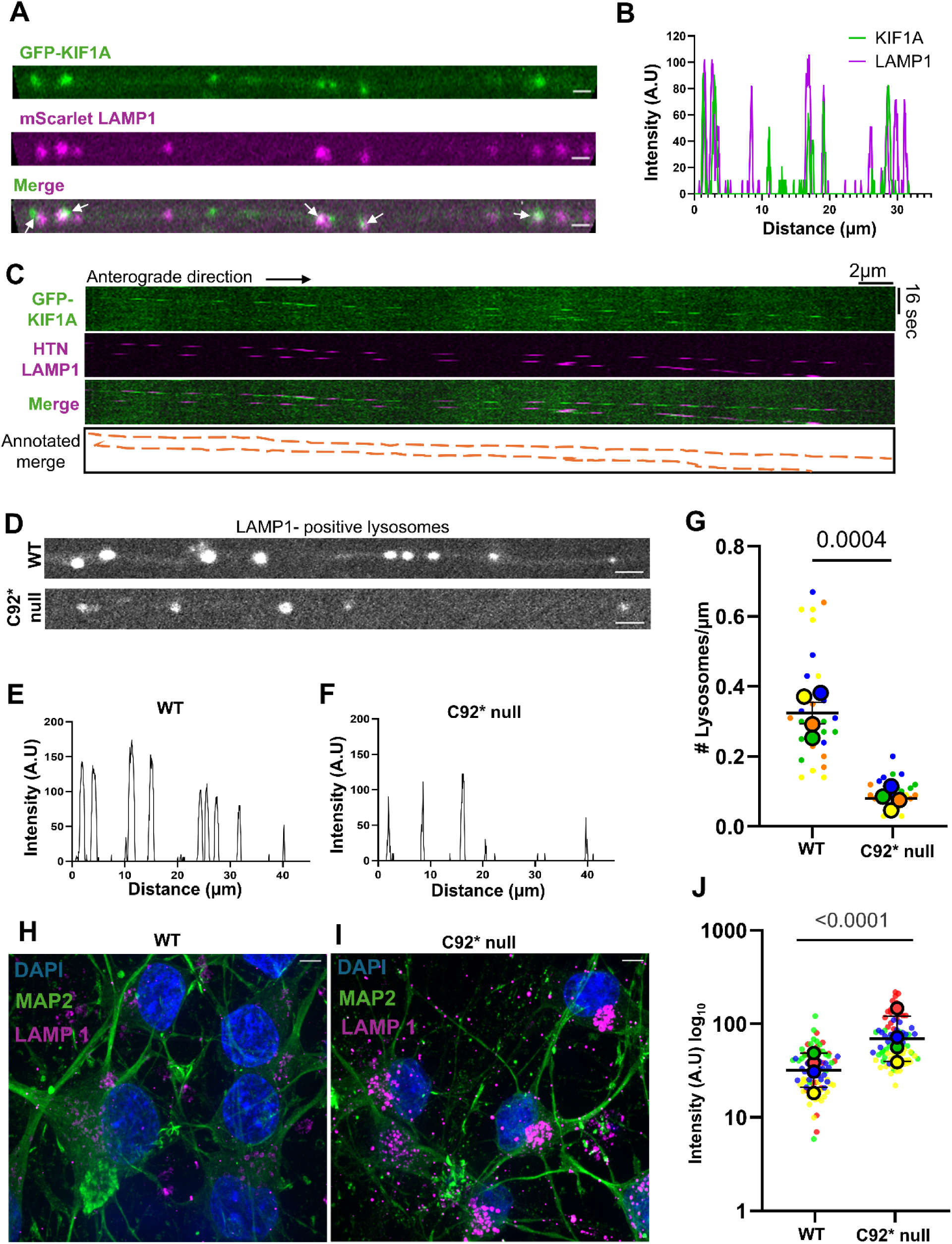
Lysosome density and localization disrupted in absence of KIF1A. **A)** Still image of axon showing KIF1A (green) puncta co-localizing with LAMP1 (magenta) puncta. Scale bars 1µm. Arrows point to colocalizing vesicles. **B)** Intensity plot showing the instances of KIF1A (green) and LAMP1 (magenta) colocalization along the axon represented in image 4A. Y-axis is intensity (mean grey value); x-axis is distance in microns. **C)** Representative kymographs with annotation showing KIF1A (green) and HTN-LAMP1 (magenta) co-trafficking events. Videos recorded at 5fps for 5 minutes. Scale bars 2µm. **D)** Representative still images of WT and C92* null axons transfected with mNeon green LAMP1. Scale bars 2µm. **E)** Intensity plot for representative WT still image (**D**). Y-axis is intensity (mean grey value); x-axis is distance in microns. **F)** Intensity plot for representative C92*null still image (**D**). Y-axis is intensity (mean grey value); x-axis is distance in microns. **G)** Superplot showing the number of lysosomes present along DIV21 WT and C92* null axons (#lysosomes/µm). Plot shows mean ± standard deviation of experimental replicates; n =28 neurons for each condition from 4 independent experiments. Small dots represent individual cells; large dots represent the average for each replicate. Each color represents 1 replicate. WT average = 0.34 vesicles/µm, C92* null average= 0.10 vesicles/µm. P-value determined by unpaired t-test. **H-I)** Representative images of WT (**H**) and C92*null (**I**) neurons showing LAMP1 positive lysosome localization in the somal region. Color-coded with DAPI (blue), Map2 (green) and LAMP1 (Magenta). Scale bars 5µm. **J)** Superplot showing the Mean grey value (intensity) of LAMP1 positive lysosomes confined to the soma in WT and C92* null neurons. Plot shows the geometric mean ± standard deviation of experimental replicates; n =85 neurons for each condition from 4 independent experiments. Small dots represent individual cells; large dots represent the average for each replicate. Each color represents 1 replicate. WT geometric mean= 33.05 a.u., 95% CI [17.42,62.69], C92* null geometric mean=69.39 a.u., 95% CI [29.53,163.1]. P-value determined by mixed- effects model.

Both kinesin-1 and KIF1A, a kinesin-3 isoform, are thought to traffic lysosomes into the axon, but our data indicate that a loss of KIF1A is sufficient to significantly decrease the density of axonal lysosomes. Previous work has suggested that kinesin-1 and kinesin-3 might have different roles in cargo transport. Evidence suggests that kinesin-1 is possibly required for trafficking through the AIS, and that kinesin-3 drives subsequent trafficking toward the distal axon.^62,64^ To test this possibility, we transfected WT and C92* null neurons with mNeon LAMP1 and imaged lysosome transport at the AIS region, defined as within 10-60µm of the soma, as well as more distally along the proximal axon (**Supplemental figure 2A**). Within the AIS, we noted similar lysosomal density between WT and C92* null neurons (**Supplemental Video 1 and 2, Supplemental figure 2B**). However, lysosome density, and especially the density of anterograde moving lysosomes, decreased in C92* null axons compared to WT distal to the AIS (**Supplemental Video 3 and 4, Supplemental figure 2C**). This suggests that KIF1A is primarily required for lysosome trafficking in the axon, distal to the AIS.

### Loss of KIF1A inhibits autophagosome maturation

To determine whether the lysosomal deficit we observed results in altered autophagosome maturation, we assessed autophagic flux using western blotting. WT and C92* null neurons were treated with DMSO or 100nM Bafilomycin A1 (BafA1) and then probed for levels of LC3 lipidation. BafA1 blocks autophagosome-lysosome fusion by neutralizing lysosomal acidification^65^ **(Figure 5A)**. WT neurons exhibit increased levels of lipidated LC3 (LC3-II) and an increased LC3-II/LC3-I ratio following treatment with BafA1 **(Supplemental figure 3A; Figure 5B-C**). In contrast, C92* null neurons exhibited a higher level of LC3-II as well as a higher LC3-II/LC3-I ratio than WT neurons at baseline, but treatment with BafA1 did not significantly increase either the level of LC3-II or the LC3-II/LC3-I ratio **(Supplemental figure 3A; Figure 5B-C**). Total LC3 levels (LC3-I+LC3-II) were unchanged across conditions (**Supplemental figure 3B**). Because the LC3-II/LC3-I ratio did not increase upon BafA1 treatment in C92* null neurons, we conclude that depletion of axonal lysosomes impairs autophagic flux. As an orthogonal assay for autophagosome maturation, we performed live-cell imaging on WT and C92* null neurons transfected with a fused GFP-mCherry-LC3 reporter. Upon autophagosome-lysosome fusion, the pH-sensitive GFP signal of the construct is quenched by the acidic environment, allowing for observation of non-acidified (GFP-mCherry-positive) and acidified (mCherry-only-positive) LC3 vesicles. In WT neurons, there were more acidified autophagosomes reaching the proximal axon (**Figure 5D & F**) accounting for about 60% of the LC3 population in this region of the cell **(Figure 5G).** In contrast, C92* null neurons showed no difference in the number of non-acidified and acidified LC3 reaching the proximal axon **(Figure 5E & F).** While about 50% of LC3-positive vesicles in the proximal axon of C92* null neurons were acidified (**Figure 5G**), there were far fewer LC3 vesicles in C92* null neurons overall, consistent with the results described in **Figure 2G-H**.

**Figure 5:**
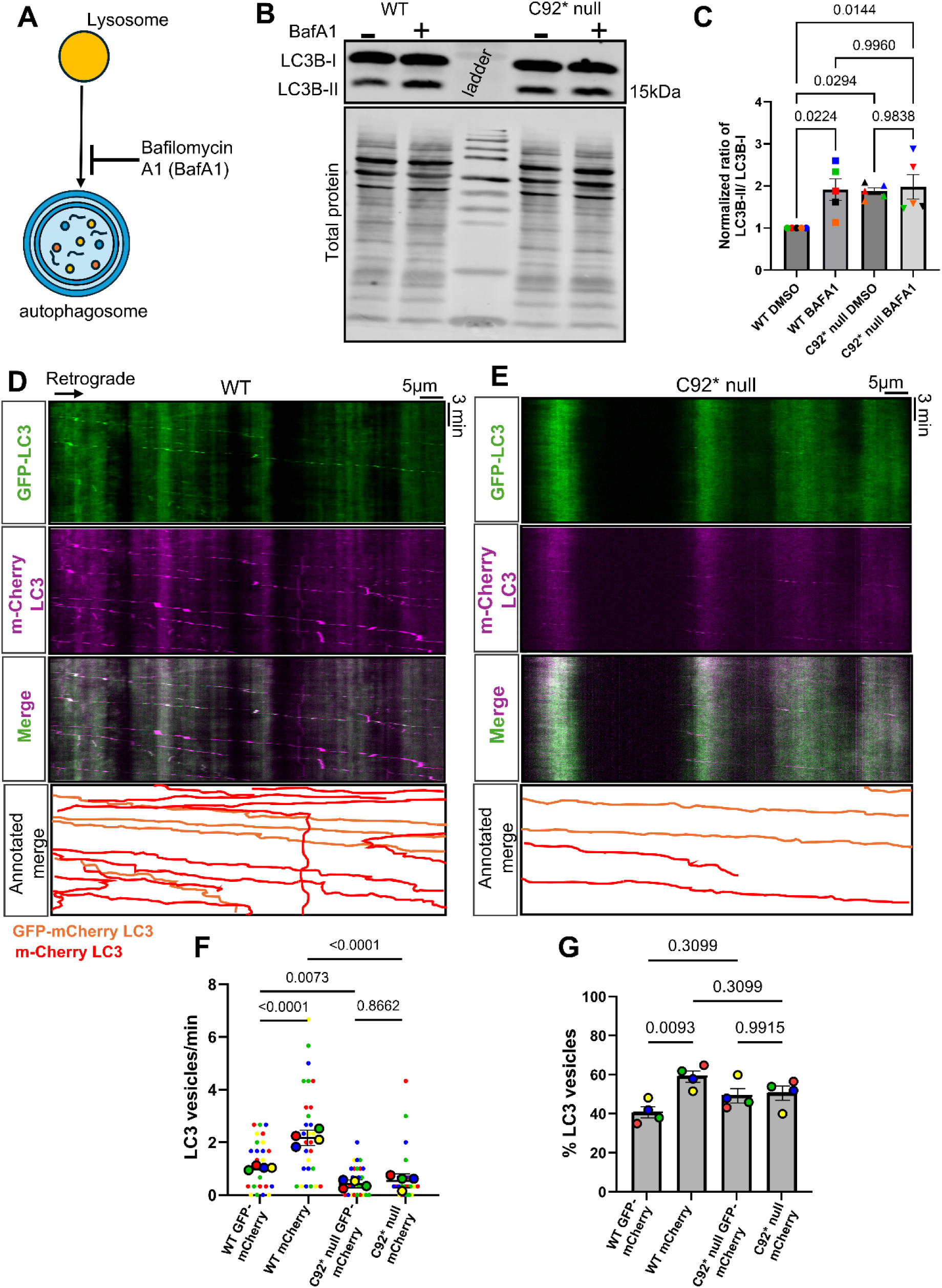
Autophagy maturation impacted by loss of KIF1A. **A)** Schematic showing the effects of Bafilomycin A (BafA1) on autophagy maturation. BafA1 blocks autophagosome-lysosome fusion by preventing lysosome acidification. **B)** Representative western blot showing WT and C92* null neurons treated with BafA1 or DMSO, then probed for LC3. **C)** Bar graph showing the ratio of LC3B-II to LC3B-I in WT and C92* null neurons treated with BafA1 or DMSO, probed for LC3 and normalized to total protein. N of 5; each color represents 1 replicate. P-values determined by One-way ANOVA with Tukey’s multiple comparison’s test. **D-E)** Representative kymographs with annotations showing GFP-mCherry dual tagged autophagosomes trafficked along WT (**D**) and C92* null proximal axons (**E**). Upon, autophagosome-lysosome fusion, the GFP is quenched resulting in two populations of LC3 positive autophagosomes: GFP-mCherry LC3 and mCherry LC3. GFP-mCherry vesicles represent autophagosomes that has not fused with a lysosome (non-acidified), and mCherry vesicles represent autophagosomes that has fused with a lysosome (acidified). Scale bars 5µm. **F)** Superplot showing the number of GFP-mCherry and mCherry LC3 trafficked along the proximal axons of WT and C92* null neurons (vesicles/min). Proximal axons defined as 100µm from the soma. Plot shows mean ± standard deviation experimental replicates; n =27 neurons for each condition from 4 independent experiments. Small dots represent individual cells; large dots represent the average for each replicate. Each color represents 1 replicate. WT GFP-mCherry LC3 average= 1.190 vesicles/min, WT mCherry LC3 average= 2.357 vesicles/min, C92* null GFP-mCherry LC3 average= 0.559 vesicles/min, C92* null mCherry LC3 average= 0.679 vesicles/min. P-values determined by One-way ANOVA with Tukey’s multiple comparison’s test. **G)** Bar graph showing the % of GFP-mCherry and mCherry LC3 trafficked along the proximal axons of WT and C92* null neurons. Plot shows mean ± standard deviation experimental replicates; n =27 neurons for each condition from 4 independent experiments. Large dots represent the average for each replicate. Each color represents 1 replicate. WT GFP-mCherry LC3 average= 40.82%, WT mCherry LC3 average= 59.18%, C92* null GFP-mCherry LC3 average=49.33%, C92* null mCherry LC3 average= 50.67%. P-values determined by One-way ANOVA with Tukey’s multiple comparison’s test.

### RNA localization decreases in axons lacking KIF1A

Recent work has identified a key role for lysosomes in the transport of RNA granules along the axon.^62,66,67,68^ Bonifacino and others have shown that blocking lysosomal transport into axons results in a depletion of mRNAs encoding mitochondrial and ribosomal proteins^63^. To investigate whether the decrease in axonal lysosomes in the C92* null neurons resulted in a similar depletion of RNA from axons, we cultured WT and C92* null neurons in microfluidic chambers and incubated with SYTO RNASelect Green Fluorescent Cell Stain to label RNA. We observed fewer SYTO-positive puncta along the axons of C92* null neurons compared to WT **(Figure 6A-B)**, though flux was not changed (**Figure 6C**). A recent study by Fenton et al., revealed that FMRP-positive RNA granules colocalize with lysosomes in neuronal processes.^69^ To investigate whether there is a deficit in the distribution of FMRP granules, we transfected WT and C92* null neurons with GFP-FMRP and quantified FMRP density along the axon. We found that there were fewer FMRP puncta distributed along C92* axons compared to WT (**Figure 6D-G**). Together, these data indicate that loss of KIF1A-mediated lysosomal trafficking leads to depletion of RNA and RNA granules from the axon.

**Figure 6:**
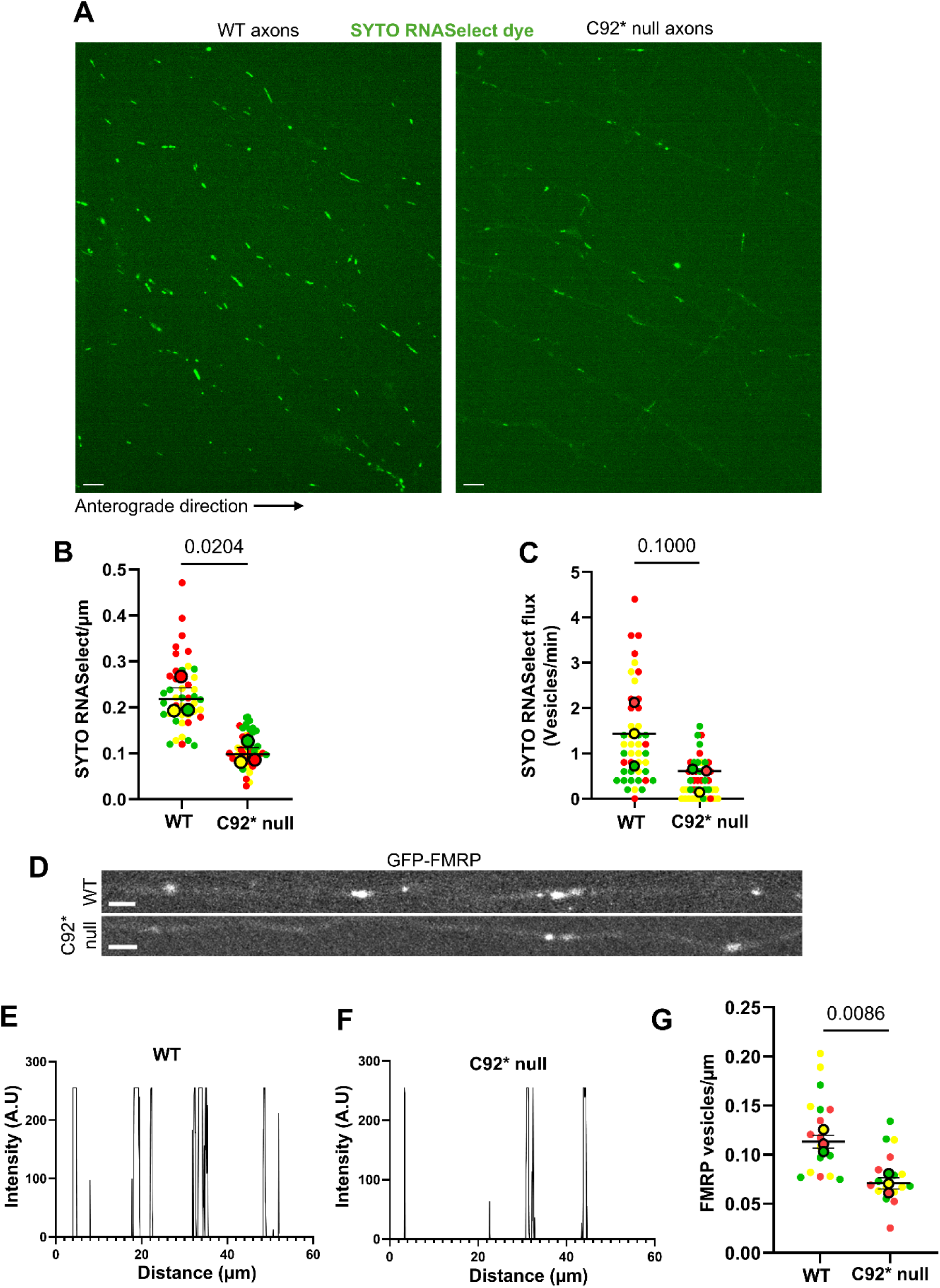
Loss of KIF1A results in less axonal RNA. **A)** Representative images showing SYTO RNASelect density in WT and C92* null neurons. WT and C92* null neurons were cultured in microfluidic chambers and incubated with SYTO RNASelect, a dye that labels RNA. Videos recorded at 1fps for 5 minutes. Scale bars 5µm. **B)** Superplot showing the number of SYTO RNASelect puncta in the axons of WT and C92* null axons (vesicle/µm). Plot shows mean ± standard deviation of experimental replicates; n =45 neurons for each condition from 3 independent experiments. Small dots represent individual cells; large dots represent the average for each replicate. Each color represents 1 replicate. WT average = 0.23 vesicles/ µm, C92* null average= 0.11 vesicles/ µm. P-value determined by unpaired t-test with Welch’s correction. **C)** Superplot showing the total vesicle flux (Vesicles/µm) of WT and C92* null axons. Plot shows median of experimental replicates; n =44-45 neurons for each condition from 3 independent experiments. Small dots represent individual cells; large dots represent the average for each replicate. Each color represents 1 replicate. WT median = 1.4 vesicles/min, C92* null median= 0.6 vesicles/min. P-value determined by Mann-Whitney test. **D)** Representative images showing GFP-FMRP puncta in WT and C92* null axons. WT and C92* null neurons were transfected with GFP-FMRP, an RNA granule. Scale bars 2 µm. **E)** Intensity plot for WT FMRP representative still image (Figure 6D). Y-axis is intensity (mean grey value); x-axis is distance in microns. **F)** Intensity plot for C92* null FMRP representative still image (Figure 6D). Y-axis is intensity (mean grey value); x-axis is distance in microns. **G)** Superplot showing the number of GFP-FMRP puncta in the axons of WT and C92* null axons (vesicle/µm). Plot shows mean ± standard deviation of experimental replicates; n =17 neurons for each condition from 3 independent experiments. Small dots represent individual cells; large dots represent the average for each replicate. Each color represents 1 replicate. WT average = 0.12 vesicles/ µm, C92* null average= 0.08 vesicles/ µm. P-value determined by unpaired t-test with Welch’s correction.

### Mitochondria dynamics are unchanged in absence of KIF1A

Defects in mitochondrial trafficking are implicated in many neurodegenerative disorders.^70^ Mitochondria transport is predominantly driven by kinesin-1,^71^ although other studies suggest that kinesin-3 is also involved in the trafficking of mitochondria along the axon.^72^ To explore this possibility, we compared mitochondrial density, flux and membrane potential in the axons of WT and C92* null neurons. We did not observe any colocalization or co-trafficking between KIF1A and mitochondria in WT neurons transfected with GFP-KIF1A and Mito-DsRed (**Figure 7A-B**). Furthermore, cells transfected with mito-Emerald green showed no difference in mitochondrial density or flux along the axon between WT and C92* null neurons (**Figure 7C-E**). However, we noticed that mitochondria along the axon of C92* null neurons consistently looked smaller and more punctate than axonal mitochondria in WT neurons. Quantification revealed that the mitochondria in C92* null neurons were shorter than mitochondria in WT neurons (**Figure 7F-G**). Next, we incubated WT and C92* neurons with TMRE, which labels functional mitochondria.^73^ Overall TMRE intensity between WT and C92* mitochondria were comparable (**Figure 7G**). Further analysis of mitochondrial health showed no difference in ATP activity (**Supplemental figure 4E**). This suggests that mitochondrial potential is not detrimentally affected by loss of KIF1A. We speculate that the observed difference in mitochondrial morphology may result from a reduction in FMRP granules in the axons of C92* null neurons, as FMRP knockout results in fragmented and dysfunctional mitochondria^69,74^

**Figure 7:**
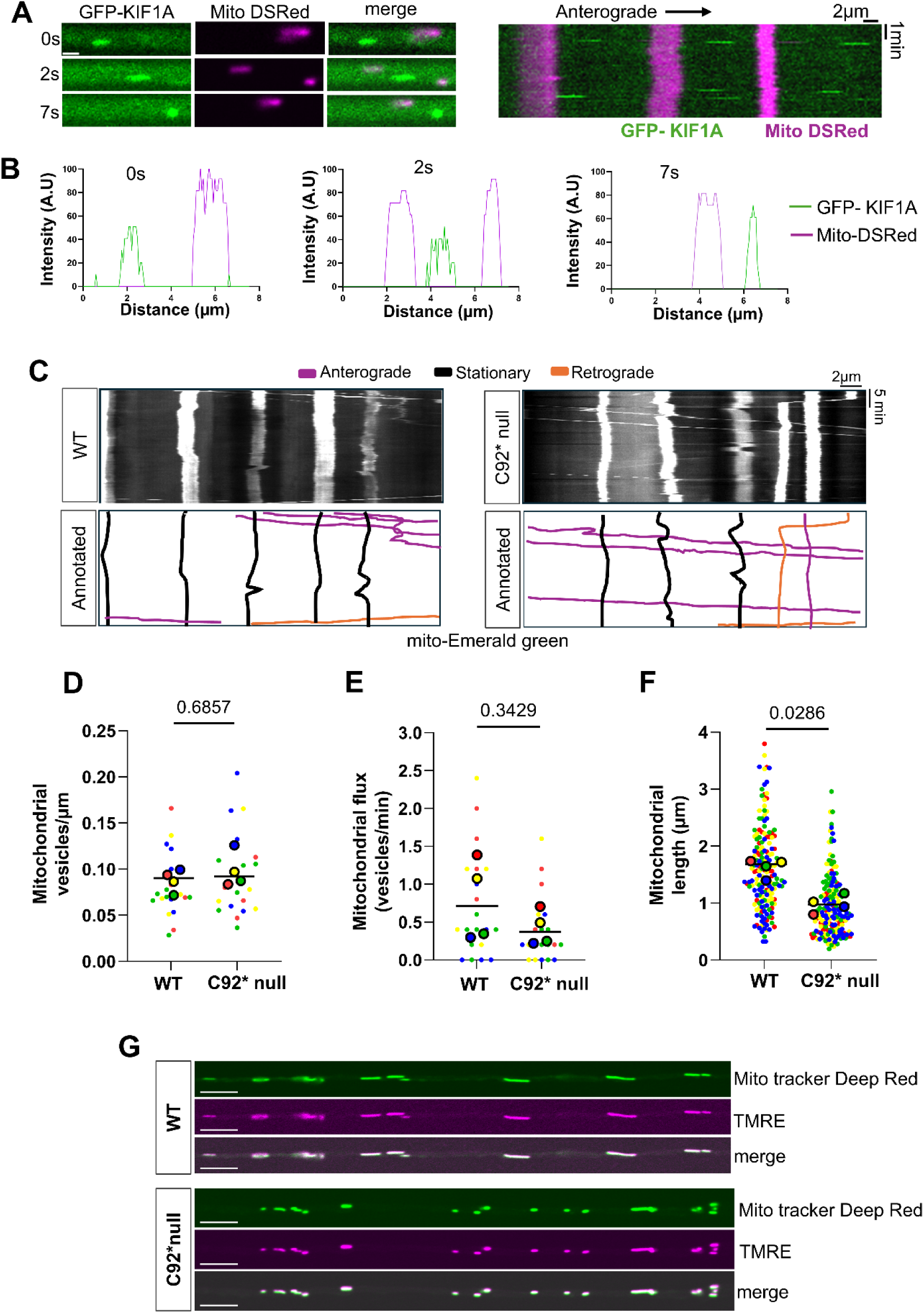
Mitochondrial dynamics unchanged in absence of KIF1A. **A)** Time series (0, 2 and 7 seconds) and kymograph showing co-movement of KIF1A and mitochondria. WT neurons were co-transfected with GFP-KIF1A (green) and mito DSRed (magenta). Scale bars 2µm. **B)** Intensity plot showing overlapping peaks of intensity between GFP-KIF1A (green) and mitoDSRed (magenta) at 0, 2 and 7 seconds. **C)** Kymographs showing WT and C92* null neuron transfected with mito-Emerald green. Videos recorded at 1fps for 5 minutes. Scale bars 2µm. **D)** Superplot showing the number of mitochondria puncta in the axons of WT and C92* null axons (vesicles/µm). Plot shows median of experimental replicates; n =20 neurons for each condition from 4 independent experiments. Small dots represent individual cells; large dots represent the average for each replicate. Each color represents 1 replicate. WT median = 0.08 vesicles/µm, C92* null median= 0.09 vesicles/ µm. P-value determined by Mann-Whitney test. **E)** Superplot showing the total mitochondria vesicle flux (Vesicles/min) of WT and C92* null axons. Plot shows median of experimental replicates; n =20 neurons for each condition from 4 independent experiments. Small dots represent individual cells; large dots represent the average for each replicate. Each color represents 1 replicate. WT median = 0.71 vesicles/min, C92* null median= 0.36 vesicles/min. P-value determined by Mann-Whitney test. **F)** Superplot showing mitochondrial length (µm) in WT and C92* null neurons. Plot shows median of experimental replicates; n =172-182 individual mitochondria for each condition from 4 independent experiments. Small dots represent individual cells; large dots represent the average for each replicate. Each color represents 1 replicate. WT median length = 1.67 µm. C92* null median length= 0.92 µm. P-value determined by Mann-Whitney test. **G)** Representative images showing Mito tracker (Green) and stained with TMRE (magenta) of WT and C92* null axons. WT and C92* null neurons were cultured in microfluidic chambers and incubated in Mito tracker and TMRE. Scale bars 5 µm.

### C92*Het neurons exhibit pronounced deficits in autophagy

Our observations establish an essential role for KIF1A in autophagosome formation and maturation in human neurons. However, KAND patients are heterozygous for the p.C92* mutation. Therefore, we explored the effects of KIF1A-mediated trafficking in autophagy comparing WT, C92* null neurons, and neurons heterozygous for C92* (C92*Het). Patients with this de novo heterozygous truncating mutation show a more severe clinical phenotype compared to other loss of function KAND mutations.^41,75^ One report highlighted a young female child heterozygous for the truncated C92* variant showing core symptoms of Rett syndrome (RTT), developmental delay, speech deficits, microcephaly and sleep disturbance.^41^ We investigated autophagy in WT, C92*Het and C92* null neurons. We first established that C92*Het neurons exhibited reduced KIF1A expression via western blot **(Supplemental figure 4A-C**). Neurons were cultured in microfluidic chambers and incubated with DAPRED to detect endogenous autophagosomes in the distal axon. We counted the density of DAPRED-positive autophagosomes 0.5-1.5µm in diameter in the distal axon, defined as within 100 µm of the axon tip. Consistent with the data described in Figure 2A-E, C92* null axons showed significantly less DAPRED puncta compared to WT axons (**Figure 8A-E).** C92*Het neurons showed increased numbers of DAPRED puncta compared to C92* null neurons, but autophagosome density was still significantly less than WT (**Figure 8A-E)**. Closer analysis of the images showed that the DAPRED puncta in C92*Het neurons were often smaller than seen in WT neurons. This led us to ask if perhaps the total number of DAPRED puncta might be the same, but due to stalled or slower membrane growth, the autophagosomes produced were smaller. We recounted DAPRED puncta in the distal axons of WT, C92*Het and C92* neurons without a size filter to get the total number. The total number of DAPRED in C92*Het and C92* null axons was still significantly lower compared to WT (**Figure 8F**). Next, we evaluated the size distribution of DAPRED puncta by comparing the proportion of larger puncta (0.5-1.5µm) to smaller puncta (<0.5 µm), within each genetic line. Highly consistent with our robust quantitative findings, this descriptive analysis revealed a progressive trend toward smaller puncta diameters in the C92*Het and the C92* null compared to WT **(Figure 8G**). We conclude that in addition to reduced total autophagosome number, both the heterozygous and homozygous C92* mutations likely impair autophagosome biogenesis, impacting both frequency and size of autophagosomes formation in the distal axons.

**Figure 8:**
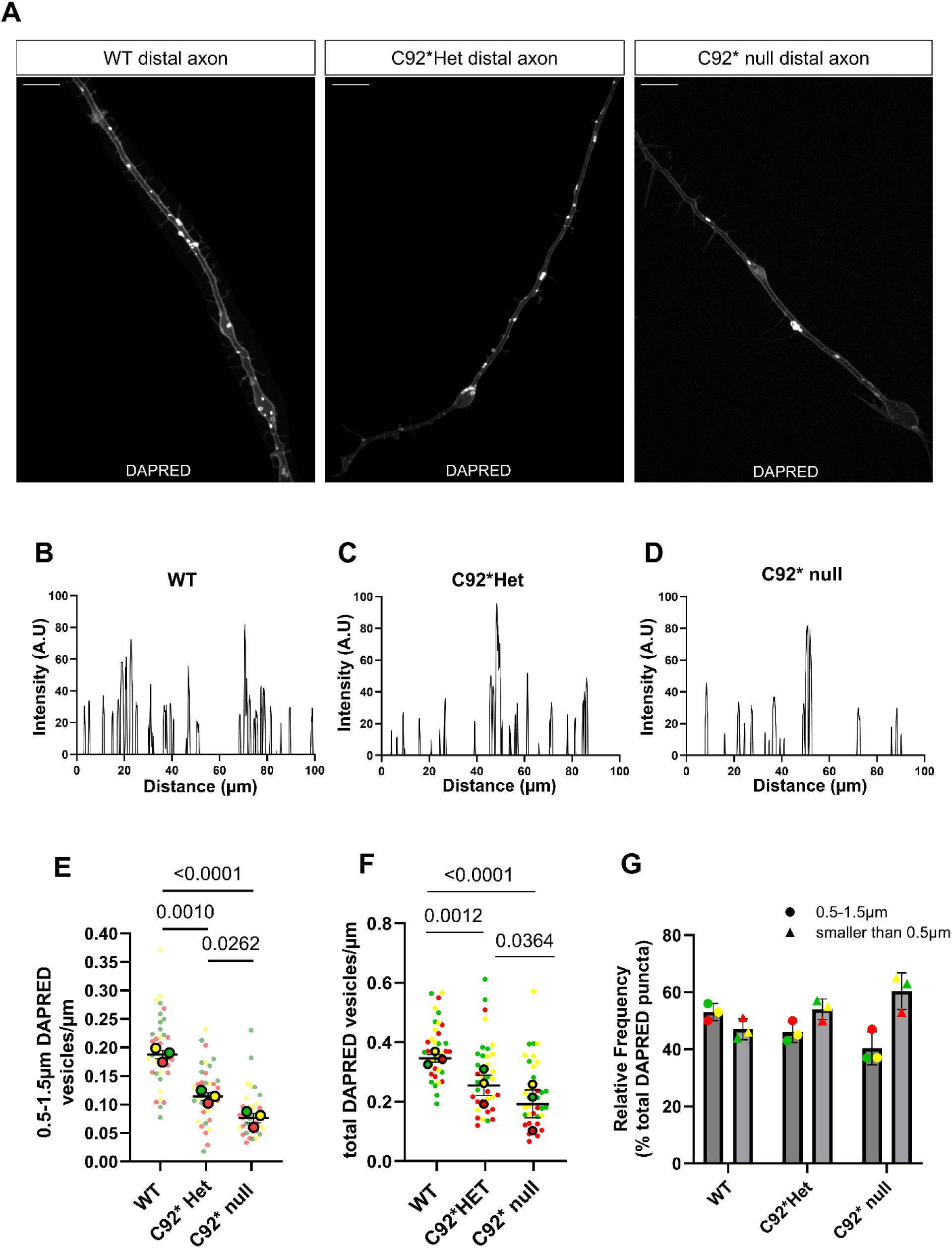
C92*Het neurons show deficits in distal autophagosome number. **A)** Representative max projection images of WT, C92*Het and C92* null distal axons. All three conditions were cultured in microfluidic chambers at DIV7 and labeled with DAPRED. Scale bars 10µm. **B)** Intensity plot for WT DAPRED representative image. Y-axis is intensity (mean grey value); x-axis is distance in microns. **C)** Intensity plot for C92*Het DAPRED representative image. Y-axis is intensity (mean grey value); x-axis is distance in microns. **D)** Intensity plot for C92* null DAPRED representative image. Y-axis is intensity (mean grey value); x-axis is distance in microns. **E)** Superplot showing the number 0.5-1.5µm diameter DAPRED puncta in the distal axons of WT, C92*Het and C92* null (vesicles/µm). Plot shows mean ± standard deviation of experimental replicates; n =36 neurons for each condition from 3 independent experiments. Small dots represent individual cells; large dots represent the average for each replicate. Each color represents 1 replicate. WT average = 0.20 vesicles/ µm, C92*Het average = 0.12, C92* null average= 0.08 vesicles/ µm. P-values determined by Ordinary one-way ANOVA with Tukey’s multiple comparison. **F)** Superplot showing the total number of DAPRED puncta in the distal axons of WT, C92*Het and C92* null axons (vesicles/µm). Plot shows mean ± standard deviation of experimental replicates; n =36 neurons for each condition from 3 independent experiments. Small dots represent individual cells; large dots represent the average for each replicate. Each color represents 1 replicate. WT average = 0.38 vesicles/ µm, C92*Het average = 0.29 vesicles/ µm, C92* null average= 0.22 vesicles/ µm. P-values determined by mixed effects model with Tukey’s multiple comparison. **G)** Size distribution profile of DAPRED puncta across WT and C92* mutants. Bar graph showing the Relative Frequency Distribution of % total DAPRED puncta within the size range (0.5-1.5µm diameter) and smaller (<0.5 µm) between WT, C92*Het, and C92* null neurons. Graph shows the mean% ± standard deviation of experimental replicates. n =36 neurons for each condition from 3 independent experiments. Each color represents 1 biological replicate. WT (0.5-1.5µm diameter) mean % = 53%; WT (<0.5 µm) a mean % = 47%. C92*Het (0.5-1.5µm diameter) mean % = 46%; C92*Het (<0.5 µm) average = 54%. C92* null (0.5-1.5µm diameter) mean % = 40%; C92* null (<0.5 µm) mean % = 60%. A descriptive, progressive trend illustrates a shift toward a higher proportion of smaller puncta from WT to C92*Het and C92* null conditions.

### Lysosome distribution and flux are altered in C92* Het neurons

Because of the striking depletion of lysosomes from the axon seen in C92* null neurons, we directly compared lysosomal density and flux in WT, C92*Het and C92* null axons transfected with mNeon-LAMP1. C92*Het axons show reduced levels of lysosome density compared to WT axons, but levels were still significantly higher than in C92* null axons (**Figure 9A-D & Figure 9F).** Furthermore, C92*Het neurons showed significantly reduced lysosomal flux compared to WT, but again significantly higher flux than C92* null neurons (**Figure 9E and 9G**). Again, western blot revealed no difference in LAMP-1 expression between WT, C92*Het and the C92* null (**Supplemental figure 4B and 4D**). Our results indicate that although the C92*Het exhibits more robust lysosomal trafficking than the complete null, the heterozygous mutation does not support WT levels of lysosomal density and lysosomal trafficking.

**Figure 9:**
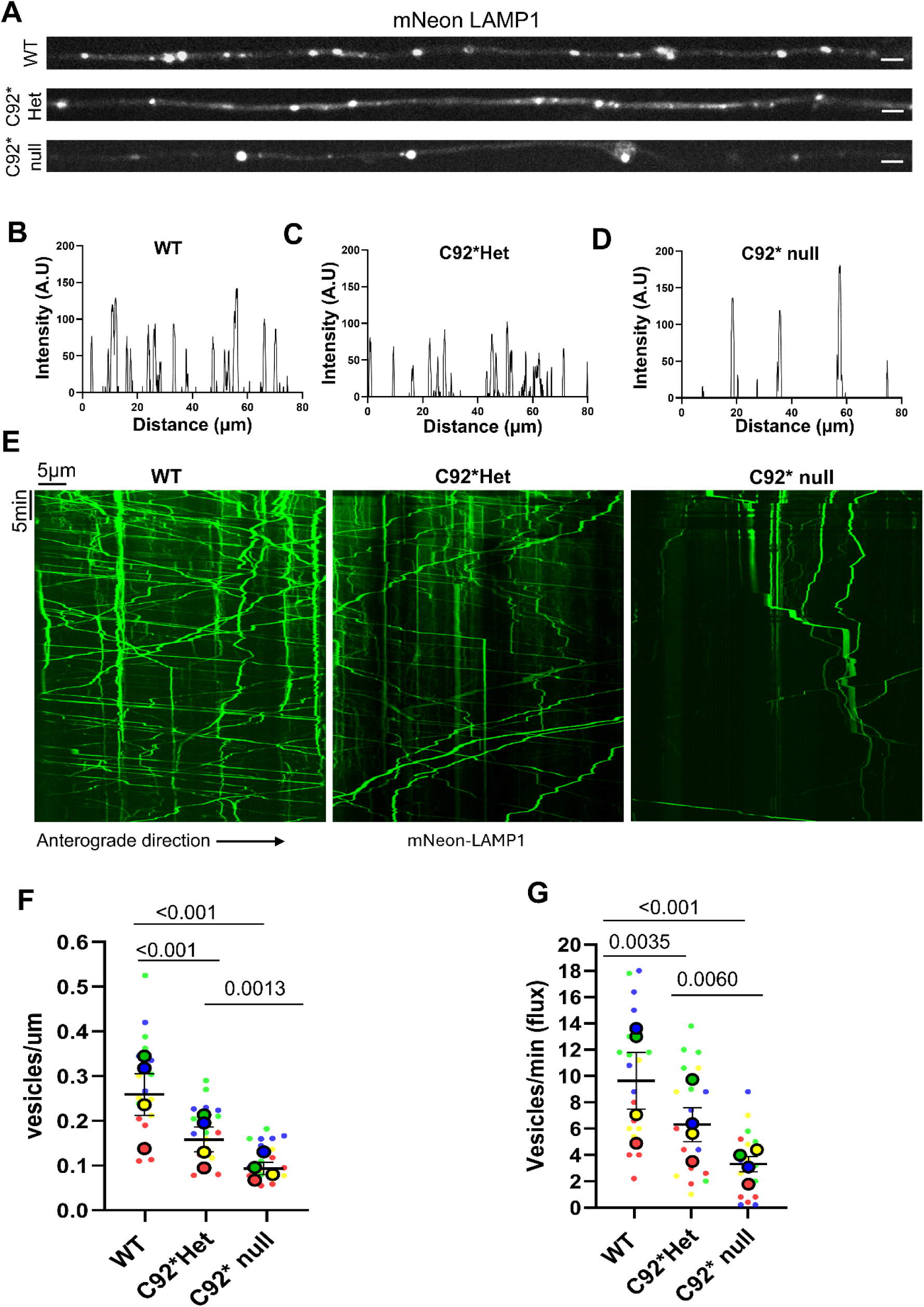
C92*Het neurons show deficits in lysosome density and flux. **A)** Representative still images of DIV21 WT, C92*Het and C92* null axons. All three conditions were transfected with mNeon green LAMP1. Scale bars 2µm. **B)** Intensity plot for representative WT still image. Y-axis is intensity (mean grey value); x-axis is distance in microns. **C)** Intensity plot for representative C92*Het still image. Y-axis is intensity (mean grey value); x-axis is distance in microns. **D)** Intensity plot for representative C92* null still image. Y-axis is intensity (mean grey value); x-axis is distance in microns. **E)** Representative kymographs of WT, C92*Het and C92* null neuron transfected with mNeon green LAMP1. Videos recorded at 5fps for 5 minutes. Scale bars 5µm. **F)** Superplot showing the number mNeon LAMP1 puncta in WT, C92*Het and C92* null axons (vesicles/µm). Plot shows mean ± standard deviation of experimental replicates; n =20 neurons for each condition from 4 independent experiments. Small dots represent individual cells; large dots represent the average for each replicate. Each color represents 1 replicate. WT average = 0.28 vesicles/ µm, C92*Het average = 0.17 vesicles/ µm, C92* null average= 0.11 vesicles/ µm. P-values determined by mixed effects model with Tukey’s multiple comparison. **G)** Superplot showing mNeon LAMP1 flux in WT, C92*Het and C92* null axons (vesicles/min). Plot shows mean ± standard deviation of experimental replicates; n =20 neurons for each condition from 4 independent experiments. Small dots represent individual cells; large dots represent the average for each replicate. Each color represents 1 replicate. WT average = 9.77 vesicles/min, C92*Het average = 6.40 vesicles/min, C92* null average= 3.35 vesicles/min. P-values determined by mixed effects model with Tukey’s multiple comparison.

### Autophagosome-lysosome fusion is reduced in C92*Het neurons

To determine whether the reduced number of both lysosomes and autophagosomes observed in p.C92*Het neurons impacts autophagosome-lysosome encounters along the axon, WT, C92*Het and C92* neurons were cultured in microfluidic chambers and incubated with lysotracker green and DAPRED to measure the densities of acidified autophagosomes along the axon. Still images for each condition were taken from the axonal compartment of the microfluidic chamber, and the number of colocalizing lysotracker green and DAPRED puncta were counted. The number of acidified autophagosomes were reduced in the C92*Het and C92* null axons compared to WT (**Figure 10 A-E**). Interestingly, the number of acidified autophagosomes was not different between C92*Het and C92* null neurons (**Figure 10E**). While about 70% of the autophagosome population in the WT axons fused with a lysosome on average **(Figure 10F)**, only 40% of the autophagosome population in the C92*Het was acidified, followed by the C92* null at 30% **(Figure 10F)**. WT axons showed a significantly higher fraction of autophagosomes colocalizing with lysosomes than seen in either C92*Het or C92* null neurons, but there was no significant difference in the fraction of autophagosomes colocalizing with lysosomes between C92*Het and C92* neurons **(Figure 10F)**. Thus, heterozygous expression of KIF1A is not sufficient to rescue autophagosome or lysosome trafficking in axons.

**Figure 10:**
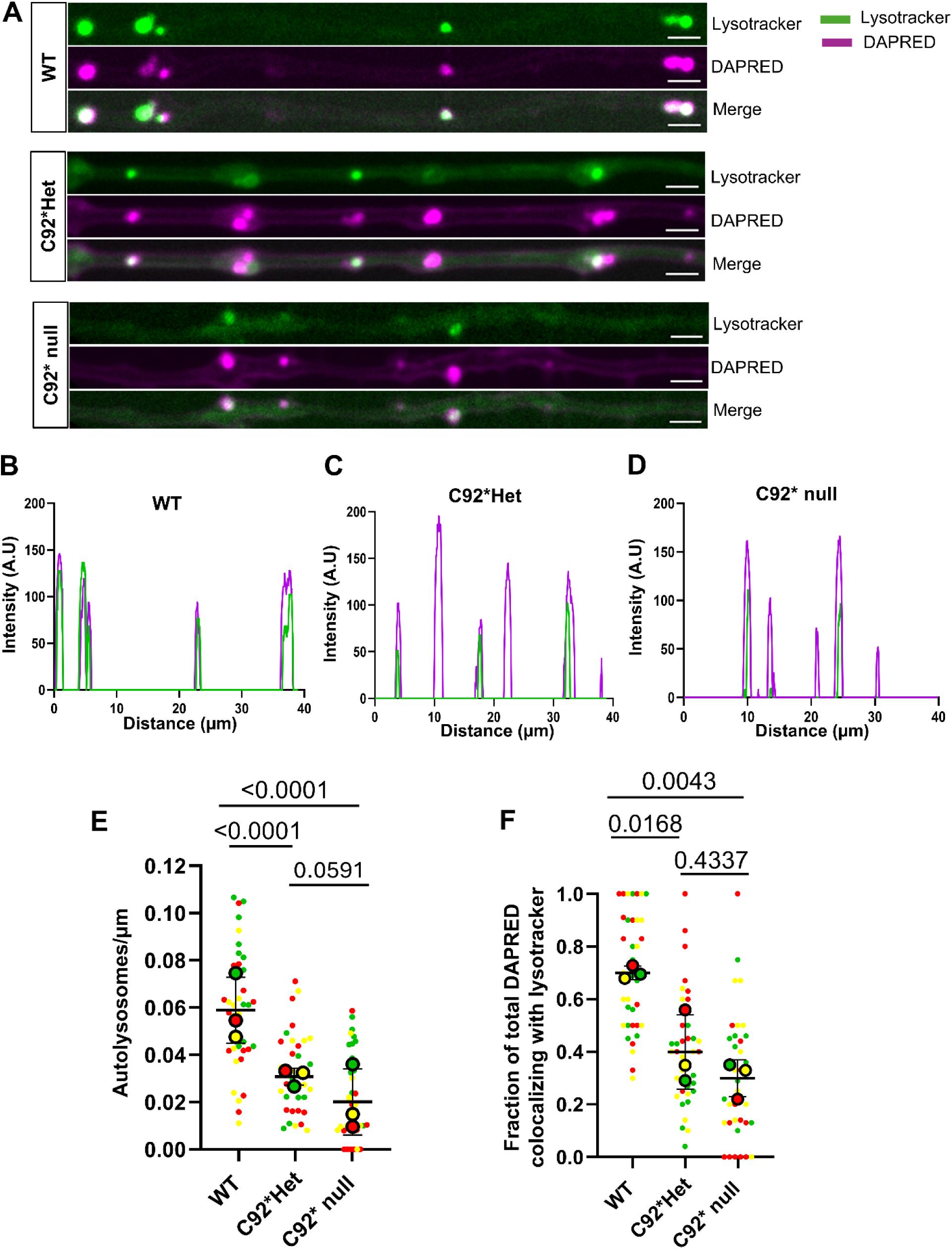
KAND mutations reduce number of axonal autophagosome-lysosome encounters. **A)** Representative still images of WT, C92*Het and C92* null axon showing lysotracker (green) and DAPRED (magenta) encounters. **B)** Intensity plot for WT representative image showing lysotracker (green) and DAPRED (magenta) encounters. Y-axis is intensity (mean grey value); x-axis is distance in microns. **C)** Intensity plot for C92*Het representative image showing lysotracker (green) and DAPRED (magenta) encounters. Y-axis is intensity (mean grey value); x-axis is distance in microns. **D)** Intensity plot for C92* null representative image showing lysotracker (green) and DAPRED (magenta) encounters. Y-axis is intensity (mean grey value); x-axis is distance in microns. **E)** Superplot showing the total number of autolysosomes (autophagosome-lysosome encounters) in the axons of WT, C92*Het and C92* null axons (autolysosomes/µm). Plot shows mean ± standard deviation of experimental replicates; n =37 neurons for each condition from 3 independent experiments. Small dots represent individual cells; large dots represent the average for each replicate. Each color represents 1 replicate. WT average = 0.059 autolysosomes/ µm, C92*Het average = 0.031 autolysosomes/ µm, C92* null average= 0.021 autolysosomes/ µm. P-values determined by mixed effects model with Tukey’s multiple comparison. **F)** Superplot showing the fraction of total autophagosomes colocalizing with lysotracker (DAPRED+Lysotracker/total DAPRED) in the axons of WT, C92*Het and C92* null axons. Plot shows mean ± standard deviation of experimental replicates; n =37 neurons for each condition from 3 independent experiments. Small dots represent individual cells; large dots represent the average for each replicate. Each color represents 1 replicate. WT average = 0.70, C92*Het average = 0.30, C92* null average= 0.20. P-values determined by Ordinary One-Way ANOVA with Tukey’s multiple comparison.

## DISCUSSION

While KAND has been primarily associated with synaptic dysfunction, the heterogeneity observed across patients and the progressive worsening of symptoms experienced by KAND patients suggest that pathways beyond synaptic function may be compromised. The autophagy-lysosomal pathway is indispensable for the health of the neuron and defects in this pathway are found across several developmental and neurodegenerative diseases. Multiple studies suggest KIF1A-mediated trafficking impacts autophagy. In C. elegans, KIF1A is shown to transport ATG9-containing vesicles.^30^ ATG9 vesicles are required for autophagosome biogenesis, facilitating lipid transfer to the growing autophagosome.^31^ Furthermore, KIF1A drives the trafficking of lysosomes along the axon. Lysosomes are the primary degradative hub in cells and required for autophagosome maturation.^76,77,78^ While these findings predict that mutations in KIF1A may detrimentally affect axonal as well as synaptic biology, this possibility has not been extensively studied in human neurons. While a recent study looked at the effects of two C-terminal missense mutations in ALS patient-derived motor neurons and found that both mutations exhibited impaired autophagy and reduced cell adhesion,^79^ we focused instead on a loss-of-function allele, C92*, as we have shown this mutation leads to a loss of KIF1A expression likely due to nonsense-mediated decay.^25^ Our findings, in conjunction with previous work, establish the importance of KIF1A in autophagy in human neurons. Specifically, loss of KIF1A results in reduced autophagosome biogenesis and reduced maturation but also induces deficits in lysosomal trafficking and RNA localization along the axon. Furthermore, our study demonstrates that even neurons heterozygous for the truncating mutation (p.C92*Het) display significant deficits in autophagy and lysosome trafficking, indicating that KIF1A is not haplo-sufficient in human neurons.

ATG9 is needed for phagophore induction and expansion, acting as a lipid scramblase. ATG9-containing vesicles originate from the trans-Golgi network (TGN), where their export is mediated by AP-4 complex.^49^ Consistent with this, loss of AP-4 caused mislocalization of ATG9 to the cell’s periphery and increased accumulation of ATG9 at the TGN.^49^ In C. elegans neurons, ATG9 vesicles are trafficked to the site of autophagosome biogenesis by the KIF1A motor;^30^ our work shows that the role of KIF1A in ATG9 trafficking is conserved in human neurons. In neurons lacking KIF1A, we observed reduced levels of axonal ATG9, but not complete loss. This suggests that there may be partial compensation by another member of the kinesin superfamily. In non-neuronal cells, the adaptor protein RUSC2 recruits KIF5B to ATG9 containing vesicles.^80^ KIF5B is ubiquitously expressed in neurons and may partially compensate for the loss of KIF1A in ATG9 trafficking, although this partial compensation is insufficient to maintain normal levels of autophagy. Interestingly, only a few ATG9 vesicles are needed to mediate formation of an autophagosome.^81^ This finding is consistent with our observations on autophagosome biogenesis; we saw slower rates of autophagy biogenesis and a reduced number of DAPRED-positive vesicles along the axon, but not complete loss. In addition to the reduced number of DAPRED-positive vesicles, we noticed that many DAPRED-positive vesicles were smaller in C92* null neurons as compared to WT. We speculate that reduced levels of ATG9 resulted in less efficient lipid transfer from the ER to growing phagophores; loss of ATG9’s scramblase activity means that lipids coming in from ATG2 cannot be properly inserted into the membrane, leading to smaller, stalled autophagosomes. ATG9 also contributes to phagophore expansion by mobilizing lipids from lipid droplets to the phagophore and mitochondria.^82^ In HeLa cells and C. elegans, ATG9 knockout led to increased size and number of lipid droplets because fatty acid transfer from lipid droplets was blocked.^82^ It will be of interest to investigate whether a similar reduction in lipid droplet mobilization can be seen in KIF1A-null neurons.

We also noted profound deficits in the axonal trafficking of lysosomes in KIF1A-null neurons, and a parallel increase in somal retention, which is consistent with prior studies.^28,79,62^ Both kinesin-3 and kinesin-1 motors are implicated in lysosomal transport. While Farías et al. (2017)^61^ concluded that kinesin-1 is the main driver of lysosomal motility in hippocampal neurons, we propose that both kinesins work together to drive the outward movement of lysosomes. Consistent with this, both kinesin-1 and kinesin-3 are regulated by the BORC complex.^29,63^ While both KIF1A and kinesin-1 drive lysosomes in the anterograde direction, we hypothesize that these motors likely act in a relay or cooperative capacity. Kinesin-1 is slower in speed but generates more force and moves preferentially along stable, acetylated or detyrosinated microtubules. In contrast, the kinesin-3 motor KIF1A is faster, generates less force, and reattaches to the microtubule more quickly than kinesin-1. The unique motility and binding properties of each kinesin likely to contribute to efficient trafficking of lysosomes along the axon. Specifically, live imaging of lysosome dynamics in WT and C92* neurons indicate similar lysosomal density proximal to the AIS, defined in these experiments as the first 10-60µm of the axon. However, in the region just distal to the AIS, we saw a clear reduction in the density of anterograde LAMP1-positive organelles in in KIF1A-null neurons. Therefore, we speculate that KIF1A assists in the axonal entry of LAMP1-positive endosomes and lysosomes by navigating the AIS. The AIS region acts as a filter between the somal and axonal compartments. Gumy et al., (2017)^83^ noted that the enrichment of MAP2 in the proximal axon of DRG neurons inhibits kinesin-1 but not kinesin-3 in the trafficking of dense core vesicles (DCVs) into the axon. Therefore, based on the different properties of the two kinesins, these motors are likely to work in tandem to traffic lysosomes in neurons.

It is important to note that loss of KIF1A did not result in a complete loss of lysosomes from the axon, as was seen with a BORC knockout,^29,63^ although lysosomal density was drastically reduced in KIF1A-null neurons. Again, this may be due to partial compensation by other motors, notably KIF1Bβ. KIF1Bβ is similar in structure to KIF1A and is ubiquitously expressed in neurons. It is also considered to be a long-range motor that is regulated by the BORC complex and traffics lysosomes to the cell periphery.^29^ Autophagosomes must fuse with lysosomes to become degradative organelles. In neurons, autophagosomes generated in the distal axon fuse with lysosomes en route to the soma. This fusion is followed by the breakdown of the inner autophagosomal membrane (IAM); this breakdown is much slower in neurons than in other cell types.^34^ Successful IAM breakdown generally occurs by the time the autolysosome reaches the proximal axon. The tandem mCherry-EGFP-LC3 marker is an excellent readout for organelle maturation as the tandem fluorescent tags are localized to the inner lumen, and the pH-dependent quenching of the GFP signifies fusion IAM breakdown.^34^ Our results show that in WT human cortical-like neurons, 60% of axonal autophagosomes have fused with a lysosome and undergone IAM breakdown by the time they reach the soma. In C92* null neurons, about 50% of axonal autophagosomes are fully acidified by the time they reach the proximal axon, though the total number of autophagosomes and lysosomes were significantly less than in WT neurons (**Figure 2G-H**, **Figure 7)**. These findings suggest that the reduced density of lysosomes in the axon is sufficient to support productive fusion with the reduced number of axonal autophagosomes being trafficked to the soma. Modeling studies indicate that in wild type neurons, each autophagosome may encounter up to ∼270 lysosomes during transport from the distal to proximal axon, yet fuse with 1-2 at most.^34^ Together, these findings suggest that autophagosome-lysosome fusion is tightly regulated by SNAREs such as syntaxin-17^84^ and thus does not require high local concentrations of lysosomes to proceed efficiently.

In addition to their degradative function, axonal lysosomes are associated with RNA granules, and disruption of lysosomal trafficking significantly depletes levels of axonal mRNAs.^63,66,85^ Local translation in the axon is essential to maintain neuronal health. Depletion of axonal mRNAs by knockout of the BORC complex or loss of the RNA granule protein FMRP both induce mitochondrial dysfunction.^63,69^ Our observations on C92* null neurons support these findings, as loss of KIF1A resulted in decreased levels of RNA and decreased density of FMRP-positive RNA granules along axons. Although we did not see changes in mitochondrial density or trafficking in axons of C92* null neurons, we did observe increased mitochondrial fragmentation as compared to WT neurons. Reduced levels of FMRP granules may contribute to deregulated fission events in the mitochondria, similar to the observations of Fenton et al. (2024)^69^. However, we did not observe any significant changes in mitochondrial polarization, suggesting that mitochondrial function was not substantially altered as was seen in FMRP knockout neurons (Fenton et al., 2024)^69^, potentially because FMRP granules were depleted but not absent from C92* null neurons. This suggests that a baseline level of FMRP localization may be sufficient to maintain the health of axonal mitochondria but complete loss of FMRP disrupts mitochondrial function.

Most mutations in KIF1A arise *de novo*. These mutations, localized throughout the coding region, have been shown to induce both hyperactive and hypoactive motor function in vitro. We previously showed that neurons expressing either hyperactive or hypoactive mutations exhibit deficits in the trafficking of synaptic vesicle precursors and inhibit synaptic formation; functional synaptic deficits were also apparent in studies using multi-electrode arrays.^25^ Here we show that KIF1A loss also disrupts the autophagy pathway, which may further contribute to the synaptic deficits and progressive neuronal loss observed in KAND patients. Neurons heavily rely on autophagy for synaptic protein turnover, pruning and synaptic maturation^86^ Furthermore, robust synaptic functions rely on local protein synthesis.^87^ Lysosome transport, which is disrupted in loss of KIF1A, plays a key role in RNA transport for local translation at the synapse. It is also possible that reduced RNA seen in our work also contributes to the synaptic deficits seen in Aiken et al., (2026).^25^ Our previous work characterizing synaptic deficits utilized gene-edited iPSC neurons homozygous for KAND mutations.^25^ Here we show that neurons expressing a heterozygous truncation mutation exhibit significant deficits in both degradative and biosynthetic pathways. Of note, the C92* mutation is heterozygous in affected KAND patients and is associated with symptoms that are more severe and occur with an earlier age of onset than other cases of KIF1A haploinsufficiency, often caused by C-terminal truncations.^41^ The C92* mutation induces a significant decrease in KIF1A expression, likely due to nonsense-mediated decay, acting as a loss-of-function allele rather than a dominant negative.^25^ Our work shows that deficits in autophagy and lysosomal trafficking are less severe in p.C92*Het neurons as compared to homozygous KIF1A-null neurons but are still significantly impaired relative to WT function. Overall, our results extend the cellular phenotypes caused by depletion of KIF1A but remain consistent with the consideration of KAND as a synaptopathy, as basal autophagy in neurons preferentially initiates at presynaptic sites in vivo.^33^ Thus, it is likely that a failure to maintain organellar and protein quality control at presynaptic sites contributes to the diverse clinical presentation of KAND patients.

### Implications for therapeutic approaches for KIF1A mutations

KAND is a devastating syndrome that is challenging to treat because it encompasses a broad range of diverse and overlapping clinical manifestations, with the severity of phenotypes observed dependent on the specific variant and its location within the primary sequence of the molecular motor. Current treatments include ASO therapy, targeted to knockdown the mutant allele. Though this treatment strategy has been shown to mitigate some symptoms of KAND in an experimental therapy, deficits remain, some of which may reflect loss-of-function due to haploinsufficiency. Our work suggests that KIF1A expression is haplo-insufficient, a consideration that should be taken into account in the design of therapies for KAND targeting expression of mutant alleles.^88,89^ While KAND often presents as a neurodevelopmental disorder, it is also neurodegenerative. Neurons are particularly vulnerable to autophagy dysfunction. Here, we show that KIF1A is required for autophagy in human neurons. The development of therapies that successfully stimulate neuronal autophagy may potentially alleviate toxic buildup caused by impaired degradative function and thus slowing disease progression. Thus, from a therapeutic perspective, approaches that can ameliorate deficits in autophagy or lysophagy may provide some benefit to KAND patients and should be considered.

---

**Supplemental figure 1:**
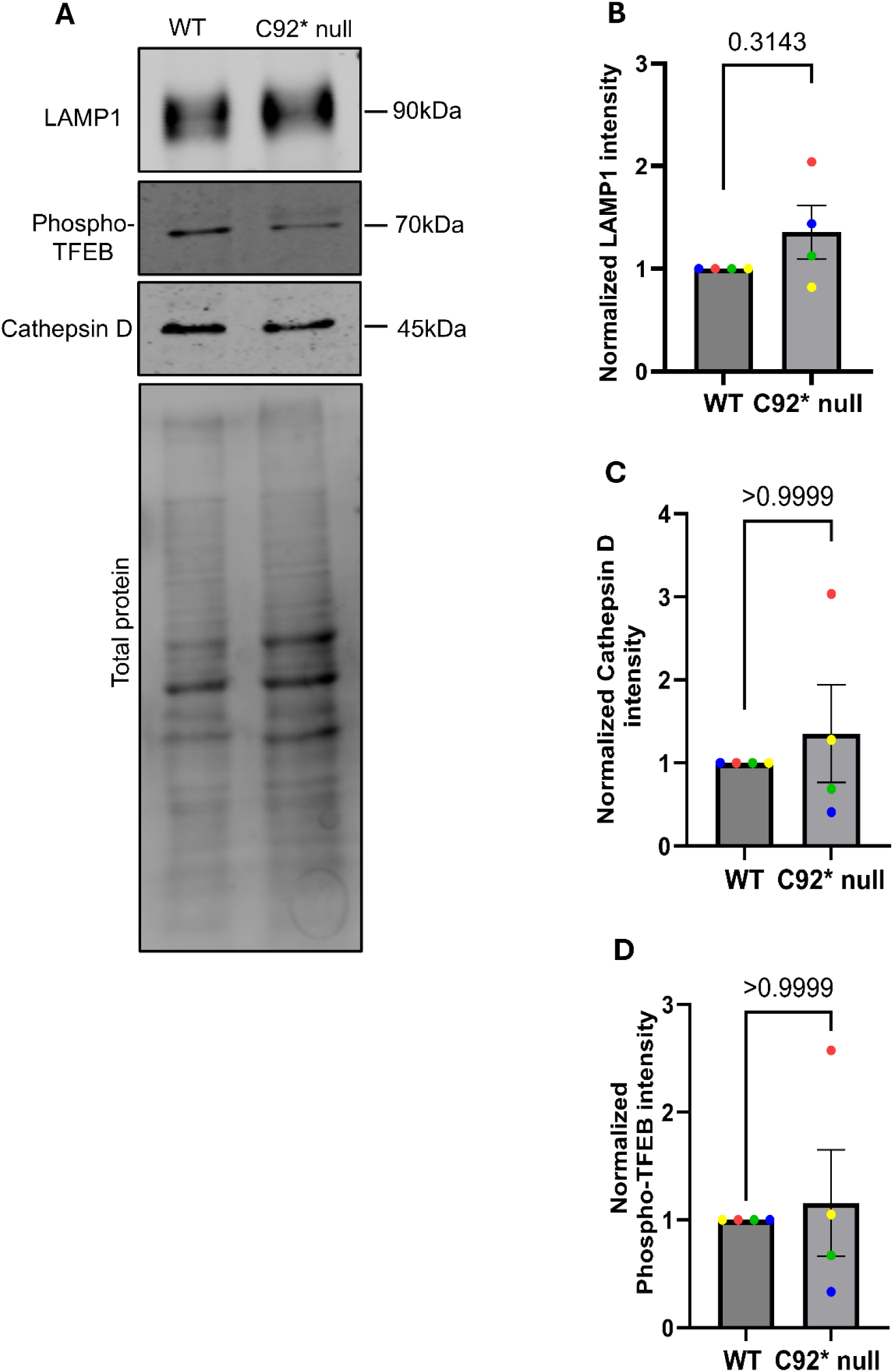
Lysosome protein expression in WT and C92* null neurons. **A)** Representative western blot image probed for LAMP1, phospo-TFEB and cathepsin D in WT and C92* null neurons. Blot was stripped each time when probing for a different protein. **B)** Bar graph showing LAMP-1 intensity normalized to total protein of WT and C92* null neurons. Graph shows mean ± standard deviation of experimental replicates (N of 4). Each color represents one replicate. P-values determined by Mann Whitney test. **C)** Bar graph showing cathepsin D intensity normalized to total protein of WT and C92* null neurons. Graph shows mean ± standard deviation of experimental replicates (N of 4). Each color represents one replicate. P-values determined by Mann Whitney test. **D)** Bar graph showing phospho-TFEB intensity normalized to total protein of WT and C92* null neurons. Graph shows mean ± standard deviation of experimental replicates (N of 4). Each color represents one replicate. P-values determined by Mann Whitney test.

**Supplemental figure 2:**
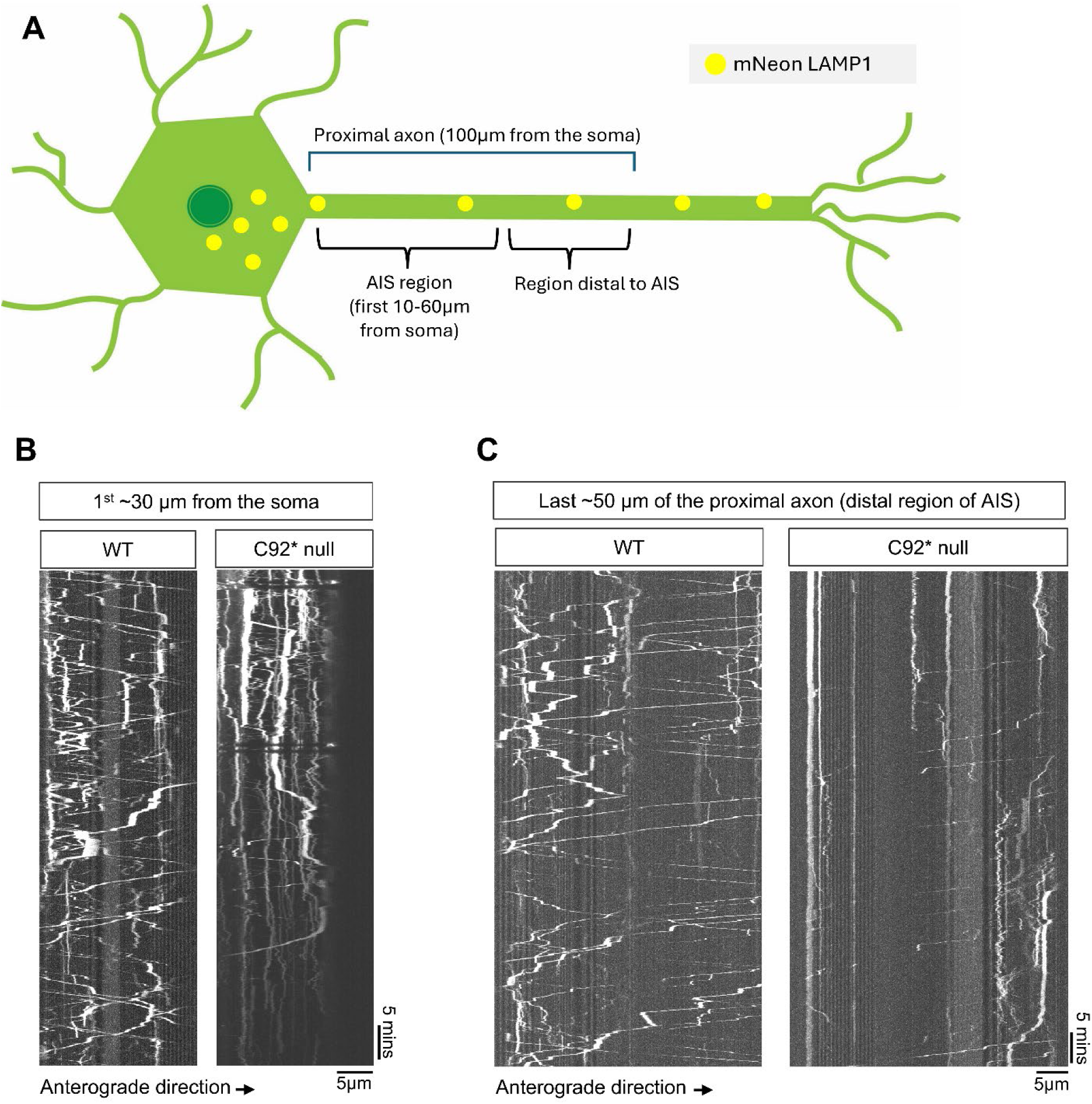
Lysosome trafficking at the proximal axon in WT and C92* null neurons. **A)** Schematic of experiment. WT and C92* null neurons were transfected with mNeon LAMP1. Videos were recorded within the AIS region (10-60µm from the soma) and after (distal to the AIS immediately following the 60µm mark). Proximal region defined as 100µm from the soma. **B)** WT Kymograph (**generated from Supplemental Video 1**) and C92* null kymograph (**generated from Supplemental Video 2**) showing LAMP1 tracks within the region of the proximal axon labeled as the AIS. 5FPS videos collected and imaged for 5 minutes. Scale bar 5µm. **C)** WT Kymograph (**generated from Supplemental Video 3**) and C92* null kymograph (**generated from Supplemental Video 4)** showing LAMP1 tracks from the same neuron (**Supplemental Video 1, Supplemental Video 2**, **Supplemental figure 2B**) but within the region of the proximal axon labeled as the region distal to the AIS. 5FPS videos collected and imaged for 5 minutes. Scale bar 5µm.

**Supplemental figure 3:**
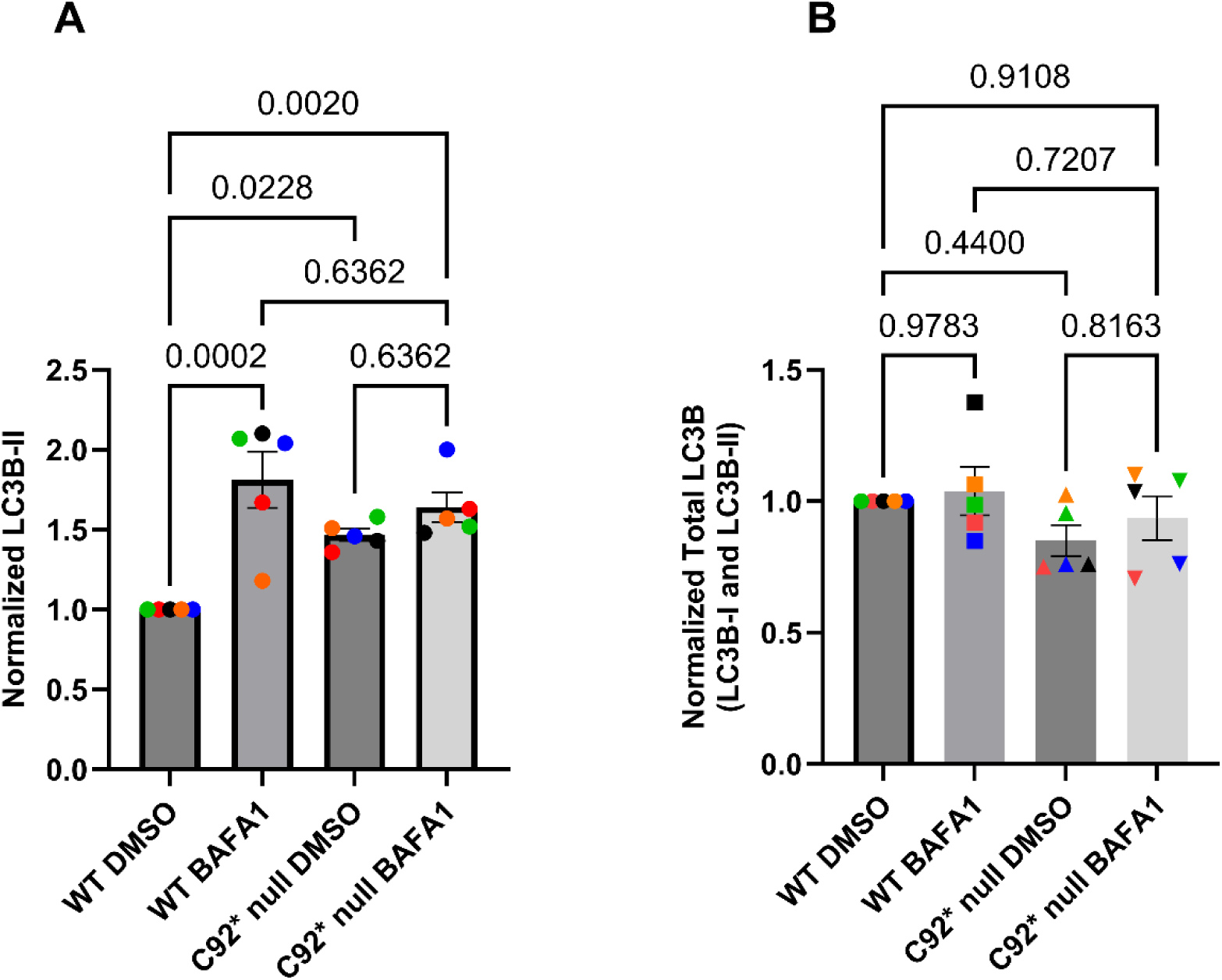
LC3B-II and total LC3B levels in WT and C92* neurons. **A)** Bar graph showing the western blot levels of LC3B-II normalized to total protein in WT and C92* neurons treated with DMSO or BafA1 (see figure 7B for western blot). Graph shows mean ± standard deviation of experimental replicates Each color represents one replicate. P-values determined by Ordinary One-way ANOVA with Tukey’s multiple comparison’s test. **B)** Bar graph showing the western blot levels of total LC3B (LC3B-I and LC3B-II) normalized to total protein in WT and C92* neurons treated with DMSO or BafA1 (see figure 7B for western blot). Graph shows mean ± standard deviation of experimental replicates (N of 5). Each color represents one replicate. P-values determined by Ordinary One-way ANOVA with Tukey’s multiple comparison’s test.

**Supplemental figure 4:**
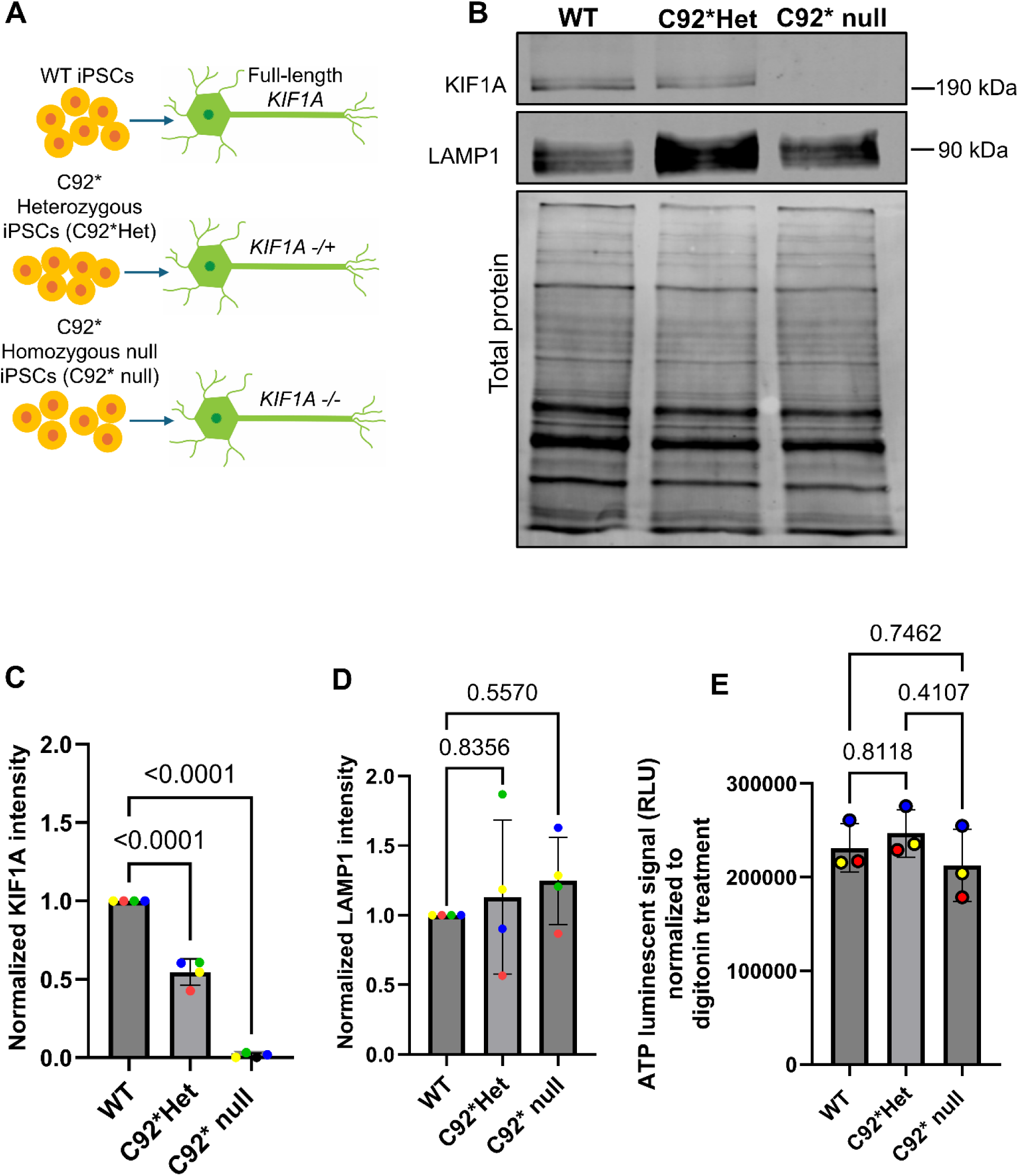
WT, C92*Het, C92* null KIF1A and LAMP1 protein expression. **A)** WT iPSCs and iPSCs gene-edited to endogenously express the C92* heterozygous truncating variant (C92*Het) and C92* null variant were differentiated to cortical-like glutamatergic neurons. **B)** Representative western blot showing KIF1A (190kDa) and LAMP1 (90kDa) protein expression for DIV21 WT, C92*Het and C92* null neurons. **C)** Quantification of relative protein levels of KIF1A DIV21 neuron lysate, corresponding to example western blot displayed in Supplemental figure 1A. Graph shows mean ± standard deviation; n = 4 independent experiments. Each color represents 1 replicate. WT average =1.00, C92*Het average = 0.55, C92* null average= 0.01. Reported *p*-values are from Ordinary one-way ANOVA with Dunnett’s multiple comparisons. **D)** Quantification of relative protein levels of LAMP-1 in DIV21 neuron lysate, corresponding to example western blot displayed in in Supplemental figure 1A. Graph shows mean ± standard deviation; N = 4 independent experiments. Each color represents 1 replicate. WT average =1.00, C92*Het average = 1.13, C92* null average= 1.23. Reported *p*-values are from Ordinary one-way ANOVA with Dunnett’s multiple comparisons. **E)** ATP assay on WT, C92*Het and C92* null with Mitochondrial ToxGlo™ Assay kit. DIV21 neurons were cultured in 96-well plates and treated with an ATP detection reagent, resulting in cell lysis and generation of a luminescent signal (RLU) proportional to the amount of ATP present. Luminescent signal normalized to digitonin treated condition. N of 3 biological replicates. WT average= 231213 RLU, C92*Het average= 246864RLU, C92* null average= 212533 RLU. P-values determined by Ordinary one-way ANOVA with Tukey’s Multiple comparison.

## Author contributions

C. B. and E.L.F.H. conceived the project and designed experiments. C.B., and J.P. performed the experiments and analyzed the data, and C.B and E.L.F.H wrote and edited the manuscript with contributions from J.P.

## Acknowledgements

We are thankful for the support of the NINDS grant R01 NS060698, Dr. Jayne Aiken, and to past and present members of the Holzbaur lab for insight and guidance on the project.

## Resource availability

Further information and requests for resources and reagents should be directed to and will be fulfilled by the lead contact, Erika Holzbaur.

### Materials availability

The gene-edited cell lines used in this study can be requested from The Jackson Laboratory.

## Methods

### Key resources table

| REAGENT or RESOURCE | SOURCE | IDENTIFIER |
| --- | --- | --- |
| **Antibodies** |  |  |
| Purified mouse monoclonal Anti-KIF1A clone 16 | BD Transduction Laboratories™ | Cat#612094 |
| monoclonal rat anti-Cathepsin D clone 204712 | Novus Biologicals | Cat# MAB1029 |
| monoclonal rabbit anti-ATG9A antibody clone | Abcam | Cat#ab108338 |
| polyclonal sheep anti-Human TGN46 | Bio-Rad | Cat# AHP500G |
| monoclonal mouse anti-MAP2 clone AP20 | Millipore Sigma | Cat#MAB3418 |
| polyclonal sheep human LAMP-1/CD107a luminal domain antibody | R&D Systems | Catalog# AF4800, RRID: AB_1026176 |
| polyclonal rabbit anti-LAMP1 antibody | Abcam | Cat# ab24170, RRID: AB_775978 |
| polyclonal rabbit LC3 antibody | Novus Biologicals | Cat# NB100-2220, RRID: AB_10003146 |
| monoclonal rabbit Phospho-TFEB (Ser211) (E9S8N) | Cell signaling Technology | Cat# 37681 |
| monoclonal mouse purified anti-Neurofilament H (NF-H), Phosphorylated antibody clone SMI 31 | Bio Legend | Cat# 801601 |
| Alexa fluor 594 donkey anti-sheep | Invitrogen | Cat# A-11016 |
| Alexa Fluor 488 goat anti-rabbit | Invitrogen | A11034 |
| Alexa fluor 633 goat anti-mouse | Invitrogen | A-21052 |
| Alexa Fluor 488 goat anti-mouse | Invitrogen | A11029 |
| IRDye 800CW Donkey anti-Rabbit | LICOR | 926-32213 |
| IRDye 800CW Donkey anti-Mouse | LICOR | 926-32212 |
| IRDye 800CW Donkey anti-Rat | LICOR | 926-32219 |
| **Bacterial and virus strains** |  |  |
| **Biological samples** |  |  |
| **Chemicals, peptides, and recombinant proteins** |  |  |
| Hoechst dye | Cell signaling technology | Cat# 40825 |
| Bafilomycin A1 READY-MADE SOLUTION | Sigma Aldrich | Cat# SML1661-.1ML |
| Mito Tracker Deep Red | Thermo Fisher Scientific | Cat# M22426 |
| SYTO RNASelect Green Fluorescent Cell Stain, 5 mM Solution in DMSO | Thermo Fisher Scientific | Cat# S32703 |
| Tetramethylrhodamine, ethyl ester (TMRE) | Thermo Fisher Scientific | Cat# T669 |
| LysoTracker™ Green DND-26, special packaging | Thermo Fisher Scientific | Cat# L7526 |
| DAPRed - Autophagy Detection (5.0 nmol) | Dojindo | Cat# D677 |
| Dimethyl sulfoxide (DMSO) | Sigma Aldrich | Cat# D2650 |
| Knockout Serum Replacement (KOSR) | Thermo Fisher Scientific | Cat# 10828010 |
| DMEM/F12 (Matrigel coating media) | Gibco | Ref# 11320-033 |
| BrainPhys without Phenol Red | StemCell | Cat# 05791 |
| BrainPhys Neuronal Medium | StemCell | Cat# 05790 |
| DMEM/F-12, HEPES (Induction media) | Gibco | Ref# 11330-032 |
| GlutaMax | Thermo Fisher Scientific | Cat# 35050061 |
| N2 Supplement | Thermo Fisher Scientific | Cat# 17502048 |
| Non-essential Amino Acids (NEAA) | Thermo Fisher Scientific | Cat# 11140050 |
| ROCK Inhibitor Y-27632 | Selleck | Cat# S1049 |
| Revert™ 700 Total Protein Stain | LICOR | Cat#926-11021 |
| NewBlot™ IR Stripping Buffer | LICOR | Cat# 928-40028 |
| EveryBlot Blocking Buffer | BioRad | Cat#12010020 |
| Halt Protease and phosphatase inhibitor | Thermo Fisher Scientific | Cat#78442 |
| Doxycycline | Sigma-Aldrich | Cat# D9891 |
| Matrigel Growth Factor Reduced | Cornig | Cat# 354230 |
| Poly-L-Ornithine (PLO) | Sigma Aldrich | Cat# P3655 |
| B27 supplement | Gibco | Cat# 17504044 |
| Laminin | Corning | Cat# 354232 |
| NT3 | PeproTech | Cat# 450-03 |
| BDNF | PeproTech | Cat# 450-02 |
| mTeSR Medium | StemCell | Cat# 85850 |
| Accutase | StemCell | Cat# 07920 |
| Fetal Bovine Serum | HyClone | Cat# SH3007103 |
| Lipofectamine Stem Transfection Reagent | Thermo Fisher Scientific | Cat# STEM00003 |
| Opti-MEM | Gibco | Ref# 31985-070 |
| ProLong Gold Antifade mountant | Life Technologies | Cat# P36930 |
| Methanol | Thermo Fisher Scientific | Cat# A545-1 |
| Transnetyx Hibernate A | Thermo Fisher Scientific | Cat# NC0442869 |
| Janelia Fluor® 646 HaloTag® Ligand | Promega | Cat# GA1121 |
| **Critical commercial assays** |  |  |
| Mitochondrial ToxGlo™ Assay | Promega | Cat#G8000 |
| Pierce™ BCA Protein Assay Kit | Thermo Fisher Scientific | Cat#23225 |
| **Deposited data** |  |  |
| **Experimental models: Cell lines** |  |  |
| Human: KOLF2.1J WT iPSCs | B. Skarnes (Jackson Laboratories, Connecticut) | RRID: CVCL_B5P3 |
| Human: KOLF2.1J C92* Homozygous null iPSCs | Jackson Laboratories | Product code JIPSC003854 |
| Human: KOLF2.1J C92* Heterozygous iPSCs | Jackson Laboratories | Product code JIPSC003852 |
| **Experimental models: Organisms/strains** |  |  |
| **Oligonucleotides** |  |  |
| **Recombinant DNA** |  |  |
| Plasmid: PGK mScarlet-LC3B | Dou et al., 2023 | Addgene #200083 (modified from Addgene #200431) |
| PGK 4xMito-mEmerald | Dou et al., 2023, | Addgene #200430 (modified from Addgene #98818) |
| Plasmid: EGFP-FMRP | gift from G. Bassell, Emory University | N/A |
| Plasmid: Halo-LAMP1 | Stavoe et al., 2019 | Addgene # 221655 |
| Plasmid: pPGK-LAMP1mNeon | Michael Ward,<br>National Institutes of Health | N/A |
| Plasmid: Halo-DFCP1 | Stavoe et al., 2019 | Modified from<br>Addgene # 38269 |
| Plasmid: mScarlet LAMP1 | This paper | Modified from<br>Addgene #1817 |
| Plasmid: pGW1: ms KIF1A FL - GFP (Internal) | Gift from Michael Silverman | N/A |
| Plasmid: Mito-DsRed | provided by. T. Schwartz, Harvard Medical School, Boston | N/A |
| Plasmid: mCherry-EGFP-LC3B | Gift from T. Johansen, University of Tromso, Norway | N/A |
| Plasmid: PB-TO-hNGN2 | Gift from iPSC Neurodegenerative Disease Initiative (iNDI) & Michael Ward | Addgene plasmid #172115 |
| Plasmid: piggyBac™ transposase vector | Transposagen | N/A |
| **Software and algorithms** |  |  |
| LI-COR Image Studio | LICOR | https://www.licor.com/bio/image-studio/<br>RRID:SCR_015795 |
| Prism version 11 | Graphpad | N/A |
| Matlab R2022a | MathWorks | https://www.mathworks.com/products/matlab.html |
| KymoSuite (custom Matlab script) | Guedes-Dias et al., 2019 | https://github.com/jnirschl/kinesin-3_guedes-dias_2018/tree/master/kymoSuite |
| KymoResliceWide | ImageJ plugin | https://imagej.net/plugins/kymoreslicewide |
| Visiview | Visitron Systems | https://www.visitron.de/ |
| RStudio: Integrated Development for R (2021.09.2 Build 382) | RStudio Team | http://www.rstudio.com/ |
| R package: nlme | Pinheiro J, Bates D, R Core Team | http://CRAN.R-project.org/package=nlme |
| Powerpoint | Microsoft | N/A |

| Other |  |  |
| --- | --- | --- |
| XonaChip® 450 µm barrier 5PK includes (1) 2 ml tube XC Pre-Coat | Xona Microfluidics | Cat# XC450 |
| Immobilon-FL PVDF membranes | Sigma | Cat# IPFL00010 |
| 35 mm #1.5 glass bottom imaging dishes | MatTek | Cat# P35G-1.5-20-C |

### Plasmids

The following plasmids were used: PGK mScarlet-LC3B (Addgene #200083);^90^ PGK 4xMito-mEmerald (Addgene #200430),^90^ EGFP–FMRP (a gift from G. Bassell, Emory University), LAMP1-HaloTag (subcloned from LAMP1-RFP into pHTC vector),^91^ pPGK-LAMP1mNeon (a gift from M. Ward, National Institutes of Health); Halo-DFCP1 (subcloned from pMXs-puro GFP-DFCP1 which was a gift from Noboru Mizushima);^91^ mScarlet LAMP1 (subcloned from LAMP1-RFP into pmScarlet -N1 vector) GFP-KIF1A (a gift from M. Silverman),Mito-DsRed (a gift from T. Schwartz, Harvard Medical School, Boston), mCherry-EGFP-LC3B (a gift from T. Johansen, University of Tromso, Norway), and PB-TO-hNGN2 (a gift from M. Ward, National Institutes of Health).

### Antibodies

The following antibodies were used for either western blots (WB) or immunofluorescence (IF): mouse monoclonal anti-KIF1A (Cat#612094; WB 1:1000), rat monoclonal anti-Cathepsin D (Cat# MAB1029; WB 1:1000), rabbit monoclonal anti-ATG9A antibody (Cat#ab108338; WB 1:1000; IF 1:200), sheep polyclonal anti-human TGN46 (Cat# AHP500G; IF 1:200), mouse monoclonal anti-MAP2 (Cat# MAB3418; IF 1:300), sheep polyclonal anti-human LAMP1 (Catalog# AF4800; WB 1:1000; IF 1:40), rabbit polyclonal anti-LAMP1 antibody (Cat# ab24170; WB 1:1000), rabbit polyclonal LC3 antibody (Cat# NB100-2220; WB 1:1000), rabbit monoclonal phospho-TFEB (Ser211) antibody (Cat# 37681; WB 1:1000), and mouse monoclonal anti-Neurofilament H (Cat# 801601; IF 1:1000). The following secondary antibodies were used: Alexa fluor 594 donkey anti-sheep (Cat# A-11016; IF 1:1000), Alexa Fluor 488 goat anti-rabbit (Cat# A-11034; IF 1:1000), Alexa fluor 633 goat anti-mouse (Cat# A-21052; IF 1:1000), Alexa Fluor 488 goat anti-mouse (Cat# A-11029; IF 1:1000), IRDye 800CW donkey anti-rabbit (Cat# 926-32213; WB 1:10,000), IRDye 800CW donkey anti-mouse (Cat# 926-32212; WB 1:10,000), and IRDye 800CW donkey anti-rat (Cat# 926-32219; WB 1:10,000).

### Cell lines

Human KOLF2.1J WT iPSCs were a gift from B. Skarnes (Jackson Laboratories, Connecticut). Human KOLF2.1J C92* Homozygous null iPSCs (JIPSC003854) and KOLF2.1J C92* Heterozygous iPSCs (JIPSC003852) were a gift from Jackson Laboratories. All three IPSC cell lines were cultured on plates coated with Growth Factor Reduced Matrigel (Corning; Cat# 354230) diluted in DMEM/F12 (Matrigel coating media, Gibco 11320-033) and fed daily with mTeSR media (Stemcell Technologies, 85850).

All three IPSC cell lines were transfected with PB-TO-hNGN2 vector (a gift from M. Ward, NIH) and transposase vector at a 2:1 ratio using lipofectamine stem (Thermo Fisher, STEM00003) and Opti-MEM (Gibco, 31985-070) to stably express doxycycline-inducible hNGN2 using a PiggyBac delivery system. After 72 hours, transfected iPSCs were selected for 48 hours with 0.5 μg/mL puromycin. Colonies stably expressing hNGN2 were frozen in cryopreservation media containing mTeSR media, Knockout Serum Replacement (KOSR, Thermo Fisher,10828010) and Dimethyl sulfoxide (DMSO) (Sigma Aldrich, D2650). All three Piggybac-delivered NGN2 IPSC lines tested negative for Mycoplasma.

### Neuronal differentiation and cell culture

All three IPSC lines stably expressing hNGN2 were thawed on Matrigel coated dishes and fed with mTeSR media. Prior to neuronal differentiation, 40-60% confluent NGN2 IPSCs were passaged with accutase (Stem Cell Technologies, 07920) and induced to a neuronal fate with doxycycline (Sigma-Aldrich, D9891) in induction media (Gibco, 11330-032) supplemented with GlutaMax (Thermo Fisher Scientific, 35050061), N2 Supplement (Thermo Fisher Scientific, 17502048) and Non-essential Amino Acids (NEAA) (Thermo Fisher Scientific, 11140050). Pre-differentiated neurons were frozen down with cryopreservation media containing BrainPhys Neuronal Medium (StemCell, 05790), Fetal Bovine Serum (HyClone, SH3007103) and DMSO. A complete protocol can be found onProtocols.io (https://doi.org/10.17504/protocols.io.e6nvwj54dlmk/v1). Pre-differentiated neurons were thawed in BrainPhys Neuronal Medium supplemented with B27 supplement (Gibco, 17504044), Laminin (Corning, 354232), NT3 (PeproTech, 450-03) and BDNF (PeproTech, 450-02) on plates coated with PLO (Sigma Aldrich, P3655) and stored in 37-degree incubators. Media changes occurred twice a week until time of experiments. Neuronal cultures day of experiment and plating density were based on the experiments to be performed (will be specified below).

### Transfection

72 hours prior to live cell imaging, DIV21 neurons cultured in 35mm glass bottom dishes (MatTek, P35G-1.5-20-C) at a density of 500k were transfected with DNA plasmid, lipofectamine stem reagent and Opti-mem and incubated at 37 degrees for 1.5 hours. For co-transfection experiments, plasmid DNA totaled from 1-3μg. A detailed protocol can be found on Protocols.io (https://doi.org/10.17504/protocols.io.x54v9dj4zg3e/

### Live cell imaging

For live cell imaging, the culture media was replaced with Transnetyx Hibernate A (Fisher Scientific, NC0442869) and supplemented with B27, NT3, Laminin and BDNF. Cells were then moved onto the microscope stage maintained at 37°C in an environmental chamber. For experiments requiring Halo-tag, 100 nM Janelia Fluor 646-Halo ligand (Promega, GA1121) was incubated with cells for 15 min followed by a 30 min washout before imaging. For DAPRED (Dojindo, D677), 1:3,000 dilution of the dye was incubated with cells for 30 minutes followed by a 2x washout prior to imaging. For Lysotracker Green (Thermo Fisher Scientific, L7526), 1:5,000 dilution was incubated with cells and imaged immediately. For MitoTracker Deep Red (Thermo Fisher Scientific, M22426), cells were incubated at 80nM for 30minutes followed by 3x washout. For SYTO RNASelect Green Fluorescent Cell Stain (Thermo Fisher Scientific, S32703), 500nM with 30-minute incubation followed by 3x washout. For TMRE (Thermo Fisher Scientific, T669), 2.5nM concentration with 30-minute incubation followed by 3x washout. Frame rate and imaging times for the experiments performed are specified in the figure legends.

Videos were acquired using an Apochromat 100x 1.49NA oil-immersion objective on an Orbital-200 CSU spinning disk confocal mounted on a Nikon Eclipse Ti stand with four laser lines (405/488/555/640) and imaged with a Hamamatsu CMOS ORCA-Fusion camera operated by a Dell workstation running VisiView Premier Image Acquisition software. Videos of the mid axon (at least 100µm from the soma or distal tip) were captured unless stated otherwise. For Xona chip experiments (Xona Microfluidics, XC450), neurons were cultured at 40k density per well and imaged at DIV7. Xonachip experiments with DAPRED and lysotracker were collected as Z-stacks at 200nm step-size.

### Immunostaining and imaging

DIV21 neurons were cultured at a density of 200k for immunostaining experiments. Cells were permeabilized with ice-cold methanol (Thermo Fisher, A545-1) for 8 min at −20°C. Cells were then washed three times with PBS and blocked for 1 h with 5% goat serum and 1% BSA in PBS. Cells were incubated overnight at 4°C with the respective primary antibodies in blocking solution. Cells were washed with PBS the next day and washed 3x with PBS. Cells were incubated with the respective secondary antibodies in blocking solution at room temperature for 1 hour. After three washes with PBS, coverslips were mounted in ProLong Gold Antifade Mountant (Life Technologies, P36930). Images were acquired as z stacks at 200 nm step-size using a PerkinElmer UltraView Vox spinning disk confocal on a Nikon Eclipse Ti Microscope. Fixed cell experiments were performed on an Apochromat 100x 1.49NA oil-immersion objective.

### Western blotting

Cell lysates were collected by washing DIV21 cells 2x with PBS prior to lysis with RIPA buffer (50mM Tris-HCl, 150mM NaCl, 0.1% Triton X-100, 0.5% sodium deoxycholate, 0.1% SDS, 2x Halt Protease and Phosphatase inhibitor (Thermo Fisher Scientific, 78442). For cells involving Bafilomycin treatment, neurons were treated with 100nM Bafilomycin A1 (Sigma Aldrich, SML1661-.1ML) overnight (∼12hours). RIPA buffer applied for 30 minutes on ice. Samples were then centrifuged, and the supernatant was collected as the lysate fraction. A BCA assay was performed on the collected lysate, and samples were denatured in sample buffer containing SDS at 95 C. Samples were resolved using SDS-PAGE gels (8% gel for KIF1A and LAMP1 protein expression; 15% gel for LC3B expression). After electrophoresis, proteins were transferred to Immobilon-FL PVDF membranes (Sigma, IPFL00010). The membrane was dried for 1hr prior to rehydration in methanol. The membranes were then stained for total protein levels, using Li-COR Revert Total Protein stain (Licor, 926-11021). The membranes were imaged using an Odyssey CLx Infrared Imaging System (Li-COR). Following imaging, membranes were destained with 0.1M NaOH supplemented with 30% Methanol. Membranes were blocked with EveryBlot Blocking Buffer (Bio-Rad, 12010020) for 15 minutes and incubated with the respective primary antibodies at 4°C overnight. Membranes were then washed 3x with 1xTBS (50mM Tris-HCl, 274mM NaCl, 9mM KCl) supplemented with 0.1% Tween 20 (TBS-T). Membranes were then incubated with the respective secondary antibodies in EveryBlot Blocking Buffer supplemented with 0.02% SDS. Membranes were incubated for 1hr and then were washed 3x with TBS-T. Membranes were then imaged, and band intensities were measured in the Li-COR Image Studio application. For membranes that were used to probe additional proteins, the membrane was stripped with 1x stripping buffer (Licor, 928-40028).

### Quantification and statistical analysis

#### Immunostaining quantification

Max projections of each channel in each image were generated. Depending on the experiment, regions were thresholded and binarized as noted. For somal ATG9 intensity, the TGN-46 channel was binarized. For ATG9 intensity at the distal axon, NFH channel was binarized. For LAMP1 somal intensity, the MAP2 signal was binarized. The binarized images were used to generate regions of interest, and then the fluorescence intensity (mean grey value) was measured within these regions of interest.

#### Kymograph generation

Kymographs were generated in FIJI using the KymoResliceWide plugin. Kymographs were manually traced using a custom MATLAB GUI (KymoSuite). Motile vesicles (FLUX) were classified as a vesicle moving with a net displacement >10 μm in the anterograde or retrograde direction. To count the density of vesicles in an axon, the first frame of the video was used to count the number of vesicles with the count particles function in FIJI. The density of vesicles was normalized to the length of the axon.

#### Autophagosome biogenesis

DIV21 neurons co-transfected with Halo-DFCP1 and mScarlet LC3 were imaged for 15 minutes at 1fps in the distal axon, defined as 100µm from the distal tip. Only LC3 puncta colocalizing with DFCP1 during the video were counted as an event. The number of colocalizing puncta were manually tracked and counted in FIJI. The number of biogenesis events were normalized to the area of the distal axon captured in the video.

#### Autophagosome density

DAPRED incubated distal axon images were collected with z-stacks and max projection images were generated in FIJI. The max projected images were trained in WEKA segmentation plugin in FIJI and count particles function with size range set to determine the number of particles. DAPRED particles were normalized to the length of the distal axon. For experiments involving DAPRED and lysotracker colocalization, z-stack images were collected in the axonal compartment of the Xona chip. Lysotracker-positive vesicles were analyzed using a single z-stack. Both channels were trained and segmented in WEKA, and FIJI’s “AND” function was used to quantify the number of colocalizing vesicles.

#### Toxglo assay

Neurons were cultured in 96-well plates in BrainPhys without Phenol red (Stem Cell, 05791) and media was partially replaced twice a week, keeping the volume of media the same across all conditions. DIV21 neurons at 100k density were treated with ATP reagent from the Mitochondrial ToxGlo™ Assay kit (Promega, G8000). The ATP luminescent signal for each condition was normalized to 80µg/ml digitonin-treated cells.

### Statistical analysis

Statistical tests were performed in Graphpad prism version 11. Experimental data was first tested for normality to determine the appropriate statistical test. Statistical tests were performed on averages of each independent replicate (N). When using averages per replicate, we graphically represented the data as superplots,^92^ plotting all data points (n) to illustrate the spread of data. P-values and specific statistical tests are reported in the figure legends. When appropriate, some statistical tests were performed in RStudio using the linear mixed effects LME; R package “nlme”). The genotype (the conditions (WT, C92* null, C92*Het) was treated as the fixed effect. The independent experiment/culture and the neuron being recorded from were treated as nested random effects, with the neuron nested within the experiment. P-values and statistical tests reported in figure legends. For all quantifications, at least three independent experiments were analyzed.

## References

1. Cason, S.E., and Holzbaur, E.L.F. (2022). Selective motor activation in organelle transport along axons. Nat Rev Mol Cell Biol 23, 699–714. 10.1038/s41580-022-00491-w.

2. Xiong, G.-J., and Sheng, Z.-H. (2024). Presynaptic perspective: Axonal transport defects in neurodevelopmental disorders. J Cell Biol 223, e202401145. 10.1083/jcb.202401145.

3. Berth, S.H., and Lloyd, T.E. (2023). Disruption of axonal transport in neurodegeneration. J Clin Invest 133, e168554. 10.1172/JCI168554.

4. Hall, D.H., and Hedgecock, E.M. (1991). Kinesin-related gene unc-104 is required for axonal transport of synaptic vesicles in C. elegans. Cell 65, 837–847. 10.1016/0092-8674(91)90391-b.

5. Lo, K.Y., Kuzmin, A., Unger, S.M., Petersen, J.D., and Silverman, M.A. (2011). KIF1A is the primary anterograde motor protein required for the axonal transport of dense-core vesicles in cultured hippocampal neurons. Neurosci Lett 491, 168–173. 10.1016/j.neulet.2011.01.018.

6. Okada, Y., Yamazaki, H., Sekine-Aizawa, Y., and Hirokawa, N. (1995). The neuron-specific kinesin superfamily protein KIF1A is a unique monomeric motor for anterograde axonal transport of synaptic vesicle precursors. Cell 81, 769–780. 10.1016/0092-8674(95)90538-3.

7. Yonekawa, Y., Harada, A., Okada, Y., Funakoshi, T., Kanai, Y., Takei, Y., Terada, S., Noda, T., and Hirokawa, N. (1998). Defect in synaptic vesicle precursor transport and neuronal cell death in KIF1A motor protein-deficient mice. J Cell Biol 141, 431–441. 10.1083/jcb.141.2.431.

8. Kondo, M., Takei, Y., and Hirokawa, N. (2012). Motor protein KIF1A is essential for hippocampal synaptogenesis and learning enhancement in an enriched environment. Neuron 73, 743–757. 10.1016/j.neuron.2011.12.020.

9. Oliver, D., Ramachandran, S., Philbrook, A., Lambert, C.M., Nguyen, K.C.Q., Hall, D.H., and Francis, M.M. (2022). Kinesin-3 mediated axonal delivery of presynaptic neurexin stabilizes dendritic spines and postsynaptic components. PLoS Genet 18, e1010016. 10.1371/journal.pgen.1010016.

10. Westerholm-Parvinen, A., Vernos, I., and Serrano, L. (2000). Kinesin subfamily UNC104 contains a FHA domain: boundaries and physicochemical characterization. FEBS Lett 486, 285–290. 10.1016/s0014-5793(00)02310-3.

11. Soppina, V., Norris, S.R., Dizaji, A.S., Kortus, M., Veatch, S., Peckham, M., and Verhey, K.J. (2014). Dimerization of mammalian kinesin-3 motors results in superprocessive motion. Proc Natl Acad Sci U S A 111, 5562–5567. 10.1073/pnas.1400759111.

12. Tomishige, M., Klopfenstein, D.R., and Vale, R.D. (2002). Conversion of Unc104/KIF1A kinesin into a processive motor after dimerization. Science 297, 2263–2267. 10.1126/science.1073386.

13. Zaniewski, T.M., and Hancock, W.O. (2023). Positive charge in the K-loop of the kinesin-3 motor KIF1A regulates superprocessivity by enhancing microtubule affinity in the one-head-bound state. J Biol Chem 299, 102818. 10.1016/j.jbc.2022.102818.

14. Benoit, M.P.M.H., Rao, L., Asenjo, A.B., Gennerich, A., and Sosa, H. (2024). Cryo-EM unveils kinesin KIF1A’s processivity mechanism and the impact of its pathogenic variant P305L. Nat Commun 15, 5530. 10.1038/s41467-024-48720-4.

15. Budaitis, B.G., Jariwala, S., Rao, L., Yue, Y., Sept, D., Verhey, K.J., and Gennerich, A. (2021). Pathogenic mutations in the kinesin-3 motor KIF1A diminish force generation and movement through allosteric mechanisms. J Cell Biol 220, e202004227. 10.1083/jcb.202004227.

16. Pyrpassopoulos, S., Gicking, A.M., Zaniewski, T.M., Hancock, W.O., and Ostap, E.M. (2023). KIF1A is kinetically tuned to be a superengaging motor under hindering loads. Proc Natl Acad Sci U S A 120, e2216903120. 10.1073/pnas.2216903120.

17. Tomaselli, P.J., Rossor, A.M., Horga, A., Laura, M., Blake, J.C., Houlden, H., and Reilly, M.M. (2017). A de novo dominant mutation in KIF1A associated with axonal neuropathy, spasticity and autism spectrum disorder. J Peripher Nerv Syst 22, 460–463. 10.1111/jns.12235.

18. Lee, J.-R., Srour, M., Kim, D., Hamdan, F.F., Lim, S.-H., Brunel-Guitton, C., Décarie, J.-C., Rossignol, E., Mitchell, G.A., Schreiber, A., et al. (2015). De novo mutations in the motor domain of KIF1A cause cognitive impairment, spastic paraparesis, axonal neuropathy, and cerebellar atrophy. Hum Mutat 36, 69–78. 10.1002/humu.22709.

19. Hamdan, F.F., Gauthier, J., Araki, Y., Lin, D.-T., Yoshizawa, Y., Higashi, K., Park, A.-R., Spiegelman, D., Dobrzeniecka, S., Piton, A., et al. (2011). Excess of de novo deleterious mutations in genes associated with glutamatergic systems in nonsyndromic intellectual disability. Am J Hum Genet 88, 306–316. 10.1016/j.ajhg.2011.02.001.

20. Esmaeeli Nieh, S., Madou, M.R.Z., Sirajuddin, M., Fregeau, B., McKnight, D., Lexa, K., Strober, J., Spaeth, C., Hallinan, B.E., Smaoui, N., et al. (2015). De novo mutations in KIF1A cause progressive encephalopathy and brain atrophy. Ann Clin Transl Neurol 2, 623–635. 10.1002/acn3.198.

21. Bernard, E., Cluse, F., Bohic, A., Hermier, M., Raoul, C., Leblanc, P., and Guissart, C. (2024). A Novel De Novo Missense Mutation in KIF1A Associated with Young-Onset Upper-Limb Amyotrophic Lateral Sclerosis. Int J Mol Sci 25, 8170. 10.3390/ijms25158170.

22. Zheng, W., He, J., Chen, L., Yu, W., Zhang, N., Liu, X., and Fan, D. (2024). Genetic link between KIF1A mutations and amyotrophic lateral sclerosis: evidence from whole-exome sequencing. Front Aging Neurosci 16, 1421841. 10.3389/fnagi.2024.1421841.

23. Sudnawa, K.K., Li, W., Calamia, S., Kanner, C.H., Bain, J.M., Abdelhakim, A.H., Geltzeiler, A., Mebane, C.M., Provenzano, F.A., Sands, T.T., et al. (2024). Heterogeneity of comprehensive clinical phenotype and longitudinal adaptive function and correlation with computational predictions of severity of missense genotypes in KIF1A-associated neurological disorder. Genet Med 26, 101169. 10.1016/j.gim.2024.101169.

24. Boyle, L., Rao, L., Kaur, S., Fan, X., Mebane, C., Hamm, L., Thornton, A., Ahrendsen, J.T., Anderson, M.P., Christodoulou, J., et al. (2021). Genotype and defects in microtubule-based motility correlate with clinical severity in KIF1A-associated neurological disorder. HGG Adv 2, 100026. 10.1016/j.xhgg.2021.100026.

25. Aiken, J., Borland, C., Marotta, N., Prosser, B.L., and Holzbaur, E.L.F. (2026). Pathogenic KIF1A variants differentially disrupt axonal trafficking and impede synaptic development. bioRxiv, 2026.01.14.699478. 10.64898/2026.01.14.699478.

26. Anazawa, Y., Kita, T., Iguchi, R., Hayashi, K., and Niwa, S. (2022). De novo mutations in KIF1A-associated neuronal disorder (KAND) dominant-negatively inhibit motor activity and axonal transport of synaptic vesicle precursors. Proc Natl Acad Sci U S A 119, e2113795119. 10.1073/pnas.2113795119.

27. Chiba, K., Takahashi, H., Chen, M., Obinata, H., Arai, S., Hashimoto, K., Oda, T., McKenney, R.J., and Niwa, S. (2019). Disease-associated mutations hyperactivate KIF1A motility and anterograde axonal transport of synaptic vesicle precursors. Proc Natl Acad Sci U S A 116, 18429–18434. 10.1073/pnas.1905690116.

28. Hummel, J.J.A., and Hoogenraad, C.C. (2021). Specific KIF1A-adaptor interactions control selective cargo recognition. J Cell Biol 220, e202105011. 10.1083/jcb.202105011.

29. Guardia, C.M., Farías, G.G., Jia, R., Pu, J., and Bonifacino, J.S. (2016). BORC Functions Upstream of Kinesins 1 and 3 to Coordinate Regional Movement of Lysosomes along Different Microtubule Tracks. Cell Rep 17, 1950–1961. 10.1016/j.celrep.2016.10.062.

30. Stavoe, A.K.H., Kargbo-Hill, S.E., Hall, D.H., and Colón-Ramos, D.A. (2016). KIF1A/UNC-104 Transports ATG-9 to Regulate Neurodevelopment and Autophagy at Synapses. Dev Cell 38, 171–185. 10.1016/j.devcel.2016.06.012.

31. Matoba, K., Kotani, T., Tsutsumi, A., Tsuji, T., Mori, T., Noshiro, D., Sugita, Y., Nomura, N., Iwata, S., Ohsumi, Y., et al. (2020). Atg9 is a lipid scramblase that mediates autophagosomal membrane expansion. Nat Struct Mol Biol 27, 1185–1193. 10.1038/s41594-020-00518-w.

32. Valverde, D.P., Yu, S., Boggavarapu, V., Kumar, N., Lees, J.A., Walz, T., Reinisch, K.M., and Melia, T.J. (2019). ATG2 transports lipids to promote autophagosome biogenesis. J Cell Biol 218, 1787–1798. 10.1083/jcb.201811139.

33. Stavoe, A.K.H., and Holzbaur, E.L.F. (2019). Axonal autophagy: Mini-review for autophagy in the CNS. Neurosci Lett 697, 17–23. 10.1016/j.neulet.2018.03.025.

34. Cason, S.E., Mogre, S.S., Holzbaur, E.L.F., and Koslover, E.F. (2022). Spatiotemporal analysis of axonal autophagosome-lysosome dynamics reveals limited fusion events and slow maturation. Mol Biol Cell 33, ar123. 10.1091/mbc.E22-03-0111.

35. Goldsmith, J., Ordureau, A., Harper, J.W., and Holzbaur, E.L.F. (2022). Brain-derived autophagosome profiling reveals the engulfment of nucleoid-enriched mitochondrial fragments by basal autophagy in neurons. Neuron 110, 967–976.e8. 10.1016/j.neuron.2021.12.029.

36. Fleming, A., Lopez, A., Rob, M., Ramakrishna, S., Park, S.J., Li, X., and Rubinsztein, D. C. (2025). How does autophagy impact neurological function? Neuroscientist 31, 349–364. 10.1177/10738584251324459.

37. Karpova, A., Hiesinger, P.R., Kuijpers, M., Albrecht, A., Kirstein, J., Andres-Alonso, M., Biermeier, A., Eickholt, B.J., Mikhaylova, M., Maglione, M., et al. (2025). Neuronal autophagy in the control of synapse function. Neuron 113, 974–990. 10.1016/j.neuron.2025.01.019.

38. Nixon, R.A., and Rubinsztein, D.C. (2024). Mechanisms of autophagy-lysosome dysfunction in neurodegenerative diseases. Nat Rev Mol Cell Biol 25, 926–946. 10.1038/s41580-024-00757-5.

39. Deng, Z., Zhou, X., Lu, J.-H., and Yue, Z. (2021). Autophagy deficiency in neurodevelopmental disorders. Cell Biosci 11, 214. 10.1186/s13578-021-00726-x.

40. Fassio, A., Falace, A., Esposito, A., Aprile, D., Guerrini, R., and Benfenati, F. (2020). Emerging Role of the Autophagy/Lysosomal Degradative Pathway in Neurodevelopmental Disorders With Epilepsy. Front Cell Neurosci 14, 39. 10.3389/fncel.2020.00039.

41. Wang, J., Zhang, Q., Chen, Y., Yu, S., Wu, X., and Bao, X. (2019). Rett and Rett-like syndrome: Expanding the genetic spectrum to KIF1A and GRIN1 gene. Mol Genet Genomic Med 7, e968. 10.1002/mgg3.968.

42. Axe, E.L., Walker, S.A., Manifava, M., Chandra, P., Roderick, H.L., Habermann, A., Griffiths, G., and Ktistakis, N.T. (2008). Autophagosome formation from membrane compartments enriched in phosphatidylinositol 3-phosphate and dynamically connected to the endoplasmic reticulum. J Cell Biol 182, 685–701. 10.1083/jcb.200803137.

43. Maday, S., Wallace, K.E., and Holzbaur, E.L.F. (2012). Autophagosomes initiate distally and mature during transport toward the cell soma in primary neurons. J Cell Biol 196, 407–417. 10.1083/jcb.201106120.

44. Maday, S., and Holzbaur, E.L.F. (2014). Autophagosome biogenesis in primary neurons follows an ordered and spatially regulated pathway. Dev Cell 30, 71–85. 10.1016/j.devcel.2014.06.001.

45. Olivas, T.J., Wu, Y., Yu, S., Luan, L., Choi, P., Guinn, E.D., Nag, S., De Camilli, P.V., Gupta, K., and Melia, T.J. (2023). ATG9 vesicles comprise the seed membrane of mammalian autophagosomes. J Cell Biol 222, e202208088. 10.1083/jcb.202208088.

46. van Vliet, A.R., Chiduza, G.N., Maslen, S.L., Pye, V.E., Joshi, D., De Tito, S., Jefferies, H.B.J., Christodoulou, E., Roustan, C., Punch, E., et al. (2022). ATG9A and ATG2A form a heteromeric complex essential for autophagosome formation. Mol Cell 82, 4324–4339.e8. 10.1016/j.molcel.2022.10.017.

47. Chumpen Ramirez, S., Gómez-Sánchez, R., Verlhac, P., Hardenberg, R., Margheritis, E., Cosentino, K., Reggiori, F., and Ungermann, C. (2023). --Atg9 interactions via its transmembrane domains are required for phagophore expansion during autophagy. Autophagy 19, 1459–1478. 10.1080/15548627.2022.2136340.

48. Matoba, K., and Noda, N.N. (2020). Secret of Atg9: lipid scramblase activity drives de novo autophagosome biogenesis. Cell Death Differ 27, 3386–3388. 10.1038/s41418-020-00663-1.

49. Mattera, R., Park, S.Y., De Pace, R., Guardia, C.M., and Bonifacino, J.S. (2017). AP-4 mediates export of ATG9A from the trans-Golgi network to promote autophagosome formation. Proc Natl Acad Sci U S A 114, E10697–E10706. 10.1073/pnas.1717327114.

50. Yang, S., Park, D., Manning, L., Kargbo-Hill, S.E., Cao, M., Xuan, Z., Gonzalez, I., Dong, Y., Clark, B., Shao, L., et al. (2022). Presynaptic autophagy is coupled to the synaptic vesicle cycle via ATG-9. Neuron 110, 824–840.e10. 10.1016/j.neuron.2021.12.031.

51. Sipos, A., Kim, K.-J., Alvarez, J.R., and Crandall, E.D. (2024). Real-Time Autophagic Flux Measurements in Live Cells Using a Novel Fluorescent Marker DAPRed. Bio Protoc 14, e4949. 10.21769/BioProtoc.4949.

52. Mizushima, N., Ohsumi, Y., and Yoshimori, T. (2002). Autophagosome formation in mammalian cells. Cell Struct Funct 27, 421–429. 10.1247/csf.27.421.

53. Mizushima, N., and Klionsky, D.J. (2007). Protein turnover via autophagy: implications for metabolism. Annu Rev Nutr 27, 19–40. 10.1146/annurev.nutr.27.061406.093749.

54. Wen, X., and Klionsky, D.J. (2016). An overview of macroautophagy in yeast. J Mol Biol 428, 1681–1699. 10.1016/j.jmb.2016.02.021.

55. Pfeifer, U. (1978). Inhibition by insulin of the formation of autophagic vacuoles in rat liver. A morphometric approach to the kinetics of intracellular degradation by autophagy. J Cell Biol 78, 152–167. 10.1083/jcb.78.1.152.

56. Mercer, T.J., Gubas, A., and Tooze, S.A. (2018). A molecular perspective of mammalian autophagosome biogenesis. J Biol Chem 293, 5386–5395. 10.1074/jbc.R117.810366.

57. Nähse, V., Stenmark, H., and Schink, K.O. (2024). Omegasomes control formation, expansion, and closure of autophagosomes. Bioessays 46, e2400038. 10.1002/bies.202400038.

58. Nähse, V., Raiborg, C., Tan, K.W., Mørk, S., Torgersen, M.L., Wenzel, E.M., Nager, M., Salo, V.T., Johansen, T., Ikonen, E., et al. (2023). ATPase activity of DFCP1 controls selective autophagy. Nat Commun 14, 4051. 10.1038/s41467-023-39641-9.

59. Jain, A., and Zoncu, R. (2026). Lysosomes as hubs of metabolic sensing and cellular homeostasis. Mol Cell 86, 533–552. 10.1016/j.molcel.2026.01.011.

60. Farfel-Becker, T., Roney, J.C., Cheng, X.-T., Li, S., Cuddy, S.R., and Sheng, Z.-H. (2019). Neuronal Soma-Derived Degradative Lysosomes Are Continuously Delivered to Distal Axons to Maintain Local Degradation Capacity. Cell Rep 28, 51–64.e4. 10.1016/j.celrep.2019.06.013.

61. Farías, G.G., Guardia, C.M., De Pace, R., Britt, D.J., and Bonifacino, J.S. (2017). BORC/kinesin-1 ensemble drives polarized transport of lysosomes into the axon. Proc Natl Acad Sci U S A 114, E2955–E2964. 10.1073/pnas.1616363114.

62. Li, C.H., Kersten, N., Özkan, N., Nguyen, D.T.M., Koppers, M., Post, H., Altelaar, M., and Farias, G.G. (2024). Spatiotemporal proteomics reveals the biosynthetic lysosomal membrane protein interactome in neurons. Nat Commun 15, 10829. 10.1038/s41467-024-55052-w.

63. De Pace, R., Ghosh, S., Ryan, V.H., Sohn, M., Jarnik, M., Rezvan Sangsari, P., Morgan, N.Y., Dale, R.K., Ward, M.E., and Bonifacino, J.S. (2024). Messenger RNA transport on lysosomal vesicles maintains axonal mitochondrial homeostasis and prevents axonal degeneration. Nat Neurosci 27, 1087–1102. 10.1038/s41593-024-01619-1.

64. Zahavi, E.E., Hummel, J.J.A., Han, Y., Bar, C., Stucchi, R., Altelaar, M., and Hoogenraad, C.C. (2021). Combined kinesin-1 and kinesin-3 activity drives axonal trafficking of TrkB receptors in Rab6 carriers. Dev Cell 56, 494–508.e7. 10.1016/j.devcel.2021.01.010.

65. Yamamoto, A., Tagawa, Y., Yoshimori, T., Moriyama, Y., Masaki, R., and Tashiro, Y. (1998). Bafilomycin A1 prevents maturation of autophagic vacuoles by inhibiting fusion between autophagosomes and lysosomes in rat hepatoma cell line, H-4-II-E cells. Cell Struct Funct *23*, 33–42. 10.1247/csf.23.33.

66. Liao, Y.-C., Fernandopulle, M.S., Wang, G., Choi, H., Hao, L., Drerup, C.M., Patel, R., Qamar, S., Nixon-Abell, J., Shen, Y., et al. (2019). RNA Granules Hitchhike on Lysosomes for Long-Distance Transport, Using Annexin A11 as a Molecular Tether. Cell *179*, 147-164.e20. 10.1016/j.cell.2019.08.050.

67. Hafner, A.-S., Donlin-Asp, P.G., Leitch, B., Herzog, E., and Schuman, E.M. (2019). Local protein synthesis is a ubiquitous feature of neuronal pre- and postsynaptic compartments. Science 364, eaau3644. 10.1126/science.aau3644.

68. Dalla Costa, I., Buchanan, C.N., Zdradzinski, M.D., Sahoo, P.K., Smith, T.P., Thames, E., Kar, A.N., and Twiss, J.L. (2021). The functional organization of axonal mRNA transport and translation. Nat Rev Neurosci 22, 77–91. 10.1038/s41583-020-00407-7.

69. Fenton, A.R., Peng, R., Bond, C., Hugelier, S., Lakadamyali, M., Chang, Y.-W., Holzbaur, E.L.F., and Jongens, T.A. (2024). FMRP regulates MFF translation to locally direct mitochondrial fission in neurons. Nat Cell Biol 26, 2061–2074. 10.1038/s41556-024-01544-2.

70. Lu, D., Feng, Y., Liu, G., Yang, Y., Ren, Y., Chen, Z., Sun, X., Guan, Y., and Wang, Z. (2023). Mitochondrial transport in neurons and evidence for its involvement in acute neurological disorders. Front Neurosci 17, 1268883. 10.3389/fnins.2023.1268883.

71. Pilling, A.D., Horiuchi, D., Lively, C.M., and Saxton, W.M. (2006). Kinesin-1 and Dynein are the primary motors for fast transport of mitochondria in Drosophila motor axons. Mol Biol Cell 17, 2057–2068. 10.1091/mbc.e05-06-0526.

72. Sharmin, R., Elkhalil, A., Pena, S., Gaddipati, P., Clark, G., Shah, P.K., Pellegrino, M.W., Shaham, S., and Ghose, P. (2025). Mitochondria transported by Kinesin-3 prevent localized calcium spiking to inhibit caspase-dependent specialized cell death. Curr Biol 35, 4932–4945.e4. 10.1016/j.cub.2025.08.065.

73. Crowley, L.C., Christensen, M.E., and Waterhouse, N.J. (2016). Measuring Mitochondrial Transmembrane Potential by TMRE Staining. Cold Spring Harb Protoc 2016. 10.1101/pdb.prot087361.

74. Shen, M., Sirois, C.L., Guo, Y., Li, M., Dong, Q., Méndez-Albelo, N.M., Gao, Y., Khullar, S., Kissel, L., Sandoval, S.O., et al. (2023). Species-specific FMRP regulation of RACK1 is critical for prenatal cortical development. Neuron 111, 3988–4005.e11. 10.1016/j.neuron.2023.09.014.

75. Paprocka, J., Jezela-Stanek, A., Śmigiel, R., Walczak, A., Mierzewska, H., Kutkowska-Kaźmierczak, A., Płoski, R., Emich-Widera, E., and Steinborn, B. (2023). Expanding the Knowledge of KIF1A-Dependent Disorders to a Group of Polish Patients. Genes (Basel) 14, 972. 10.3390/genes14050972.

76. Perera, R.M., and Zoncu, R. (2016). The Lysosome as a Regulatory Hub. Annu Rev Cell Dev Biol 32, 223–253. 10.1146/annurev-cellbio-111315-125125.

77. Mahapatra, K.K., Mishra, S.R., Behera, B.P., Patil, S., Gewirtz, D.A., and Bhutia, S.K. (2021). The lysosome as an imperative regulator of autophagy and cell death. Cell Mol Life Sci 78, 7435–7449. 10.1007/s00018-021-03988-3.

78. Roney, J.C., Cheng, X.-T., and Sheng, Z.-H. (2022). Neuronal endolysosomal transport and lysosomal functionality in maintaining axonostasis. J Cell Biol 221, e202111077. 10.1083/jcb.202111077.

79. Zhao, M., Wang, J., Liu, M., Xu, Y., Huang, J., Zhang, Y., He, J., Gu, A., Liu, M., and Liu, X. (2024). KIF1A, R1457Q, and P1688L Mutations Induce Protein Abnormal Aggregation and Autophagy Impairment in iPSC-Derived Motor Neurons. Biomedicines *12*, 1693. 10.3390/biomedicines12081693.

80. Guardia, C.M., Jain, A., Mattera, R., Friefeld, A., Li, Y., and Bonifacino, J.S. (2021). RUSC2 and WDR47 oppositely regulate kinesin-1-dependent distribution of ATG9A to the cell periphery. Mol Biol Cell 32, ar25. 10.1091/mbc.E21-06-0295.

81. Yamamoto, H., Kakuta, S., Watanabe, T.M., Kitamura, A., Sekito, T., Kondo-Kakuta, C., Ichikawa, R., Kinjo, M., and Ohsumi, Y. (2012). Atg9 vesicles are an important membrane source during early steps of autophagosome formation. J Cell Biol 198, 219–233. 10.1083/jcb.201202061.

82. Mailler, E., Guardia, C.M., Bai, X., Jarnik, M., Williamson, C.D., Li, Y., Maio, N., Golden, A., and Bonifacino, J.S. (2021). The autophagy protein ATG9A enables lipid mobilization from lipid droplets. Nat Commun 12, 6750. 10.1038/s41467-021-26999-x.

83. Gumy, L.F., Katrukha, E.A., Grigoriev, I., Jaarsma, D., Kapitein, L.C., Akhmanova, A., and Hoogenraad, C.C. (2017). MAP2 Defines a Pre-axonal Filtering Zone to Regulate KIF1-versus KIF5-Dependent Cargo Transport in Sensory Neurons. Neuron 94, 347–362.e7. 10.1016/j.neuron.2017.03.046.

84. Itakura, E., Kishi-Itakura, C., and Mizushima, N. (2012). The hairpin-type tail-anchored SNARE syntaxin 17 targets to autophagosomes for fusion with endosomes/lysosomes. Cell 151, 1256–1269. 10.1016/j.cell.2012.11.001.

85. Cioni, J.-M., Lin, J.Q., Holtermann, A.V., Koppers, M., Jakobs, M.A.H., Azizi, A., Turner-Bridger, B., Shigeoka, T., Franze, K., Harris, W.A., et al. (2019). Late Endosomes Act as mRNA Translation Platforms and Sustain Mitochondria in Axons. Cell 176, 56–72.e15. 10.1016/j.cell.2018.11.030.

86. Lu, J., Di Florio, D.N., Boya, P., Maday, S., Springer, W., and Chu, C.T. (2026). Autophagy and mitophagy at the synapse and beyond: implications for learning, memory and neurological disorders. Autophagy 22, 10–52. 10.1080/15548627.2025.2581217.

87. Rajgor, D., Welle, T.M., and Smith, K.R. (2021). The Coordination of Local Translation, Membranous Organelle Trafficking, and Synaptic Plasticity in Neurons. Front Cell Dev Biol 9, 711446. 10.3389/fcell.2021.711446.

88. Lin, Q., Verden, D., Christodoulou, J., Gold, W.A., and Kaur, S. (2025). KIF1A-associated neurological disorders: therapeutic opportunities and challenges. Eur J Hum Genet. 10.1038/s41431-025-01978-8.

89. Ziegler, A., Carroll, J., Bain, J.M., Sands, T.T., Fee, R.J., Uher, D., Kanner, C.H., Montes, J., Glass, S., Douville, J., et al. (2024). Antisense oligonucleotide therapy in an individual with KIF1A-associated neurological disorder. Nat Med 30, 2782–2786. 10.1038/s41591-024-03197-y.

90. Dou, D., Smith, E.M., Evans, C.S., Boecker, C.A., and Holzbaur, E.L.F. (2023). Regulatory imbalance between LRRK2 kinase, PPM1H phosphatase, and ARF6 GTPase disrupts the axonal transport of autophagosomes. Cell Rep *42*, 112448. 10.1016/j.celrep.2023.112448.

91. Stavoe, A.K., Gopal, P.P., Gubas, A., Tooze, S.A., and Holzbaur, E.L. (2019). Expression of WIPI2B counteracts age-related decline in autophagosome biogenesis in neurons. Elife 8, e44219. 10.7554/eLife.44219.

92. Lord, S.J., Velle, K.B., Mullins, R.D., and Fritz-Laylin, L.K. (2020). SuperPlots: Communicating reproducibility and variability in cell biology. J Cell Biol 219, e202001064. 10.1083/jcb.202001064.

