## Supplemental video legends for "KIF1A-mediated trafficking is required for neuronal autophagy in human neurons"

**Supplemental videos**

**Video S1** (Supplemental video one) = 30fps video showing a DIV21 WT neuron transfected with mNeon LAMP1 at the region defined as the AIS. Video recorded at 5fps for 5 minutes.

**Video S2** (Supplemental video two) = 30fps video showing a DIV21 C92* null neuron transfected with mNeon LAMP1 at the region defined as the AIS. Video recorded at 5fps for 5 minutes.

**Video S3** (Supplemental video three) = 30fps video showing the same DIV21 WT neuron in **Supplemental Video 1**, but at the region distal to the AIS. Text and arrow signify the anterograde direction.

**Video S4** (Supplemental Video four) = 30fps video showing the same DIV21 C92* null neuron in **Supplemental Video 2**, but at the region distal to the AIS. Text and arrow signify the anterograde direction.
